# Lactate Blocks Tertiary Lymphoid Structure Formation by Inhibiting B Cell Chemotaxis

**DOI:** 10.1101/2025.08.30.673054

**Authors:** Victoire Boulat, Elena Alberts, Paul C. Driscoll, Miu Shing Hung, Ana Cunha, Jelmar Quist, Fangfang Liu, Carin Andrea Brundin, Ananya Bhalla, Lidia Avalle, James Rosekilly, Lucy Ryan, Cheryl Gillett, Molly Strom, Valeria Poli, James I. MacRae, Anita Grigoriadis, Dinis Pedro Calado

## Abstract

Tertiary lymphoid structures (TLS) and B cell infiltration are strong predictors of immunotherapy success across cancers, including triple-negative breast cancer (TNBC). However, immune-cold TNBCs often lack both features. Here, we identify a tumor-intrinsic mechanism that actively suppresses B cell recruitment. Despite evidence of B cell responses in cancer-associated lymph nodes (cLNs), B cells fail to infiltrate TNBC tumors or form TLS. This exclusion is not simply due to chemokine deficiency as exogenous chemokine addition fails to restore B cell migration. Using fractionation and metabolic profiling, we identify lactate as a dominant tumor-secreted metabolite that directly impairs B cell chemotaxis by disrupting mitochondrial metabolism. *In vivo*, combining lactate inhibition with engineered chemokine secretion promotes cLN-derived B cell infiltration and enables TLS formation, particularly when coupled with CD40 stimulation. Transcriptomics analyses across several human cancer datasets strengthen the association between high glycolytic activity with poor B-cell infiltration in chemokine-rich tumors. Together, our findings reveal lactate as a key metabolic barrier to B cell trafficking and TLS induction, suggesting that metabolic reprogramming may provide an avenue to convert “immune-cold” tumors into TLS-rich, immunologically responsive microenvironments.

## Main

The tumor microenvironment (TME) plays a central role in both spontaneous and therapy-induced resistance to anti-tumor immunity^1^. Immune checkpoint inhibitors (ICIs), which aim to reinvigorate suppressed immune responses, have shown efficacy in a subset of patients with advanced malignancies^2^, including triple-negative breast cancer (TNBC), an aggressive subtype defined by the absence of hormone receptors and human epidermal growth factor receptor 2 (HER2)^3^. However, despite clinical benefit in some TNBC cases, response rates remain unpredictable, with most advanced TNBC cancer failing to respond^4,5^.

The abundance of tumor-infiltrating lymphocytes (TILs) is a strong prognostic marker across cancers, including TNBC^6–8^. Spatial distribution and immune cell composition within the TME also critically shape therapeutic response^9^. Tumors have therefore been categorized into three spatial immunophenotypes: “inflamed” (or “hot”), with diffuse lymphocyte infiltration; “excluded”, with immune cells restricted to the tumor margins; and “ignored” which are largely devoid of immune cells^9^. In TNBC, “inflamed” tumors are associated with better outcomes and greater responses to ICIs^10^; but most cases fall into the excluded or ignored categories i.e., “cold”, limiting therapeutic success.

CD8⁺ T cells have historically been viewed as the primary effectors of ICI therapy^11^. However, B cells are increasingly recognized as important mediators of anti-tumor immunity^12^, with strong associations to improved survival across multiple cancers^13^, including TNBC^14–16^. Their impact appears to depend on spatial localization within the TME^17^. In solid tumors, B cells are found either as dispersed infiltrates or organized into tertiary lymphoid structures (TLS), ectopic aggregates of B and T cells that form in response to chronic inflammation^17^. The presence and organization of TLS are consistently associated with improved survival and enhanced responses to ICIs across cancer types^18^, including TNBC^19–21^. In TNBC, the strongest prognostic value is observed when B cells are located intratumorally^22^ and TLS are well-organized^21^. However, such features are present in only a minority of cases^10^. The majority of TNBCs are “cold” tumors, often lacking both intratumoral B cells and TLS, posing a major barrier to effective treatment^23^.

Immune cell exclusion from tumors can result from both passive mechanisms, such as poor antigenicity^24^, stromal barriers^25^, or insufficient chemotactic signals^26,27^; as well as active mechanisms, including immunosuppressive cytokines^28^, oncogenic signaling^29,30^, and immunosuppressive cells^31^ or factors^32^ in the TME. While much of our understanding comes from studies on T cells^33^, B cell exclusion appears even more pronounced^11^, suggesting distinct regulatory barriers^34^. Given the strong link between B cells, TLS, and immunotherapy response, uncovering how B cell trafficking is suppressed could open new avenues to convert “cold” tumors into “hot” ones. However, progress has been limited by the lack of preclinical models that exhibit TLS formation^35,36^.

In this study, we uncover a tumor-intrinsic mechanism that actively excludes B cells from the primary site. Although chemokine expression correlates with B cell infiltration in patients, we show that tumor-derived lactate disrupts murine B cell responsiveness to chemotactic signals, blocking their migration from cancer-associated lymph nodes (cLNs) to the tumor. This metabolic barrier overrides chemokine-driven recruitment and TLS formation in a mouse model of TNBC. Patient data further reveal that B cell infiltration is highest in tumors with high chemokine expression and low glycolytic activity. Together, our findings position lactate as a dominant suppressor of B cell trafficking and support a therapeutic strategy that combines metabolic inhibition with chemokine delivery to promote TLS formation and convert immune-cold tumors into inflamed, immunotherapy-responsive sites.

### TNBC tumors exclude B cells originating from lymph nodes

To investigate the basis of B cell exclusion in TNBC, we first profiled the immune landscape of a well-characterized “immune cold” orthotopic TNBC mouse model (**Figure 1A**). Fourteen days after orthotopic 67NR^37^ tumor cell implantation, flow cytometry revealed that B cells were the least abundant tumor-infiltrating lymphocyte population, with T cells and macrophages being approximately 10-fold and 60-fold more abundant, respectively (**Figure 1B-C**). Immunofluorescence confirmed their scarcity within the tumor bed, with only sparse and occasional peritumoral clustering (**Figure 1D**). These findings suggest that murine TNBC tumors present a profoundly non-permissive environment for B cell infiltration, closely mirroring immune-cold TNBC lesions in patients.

**Figure 1.**
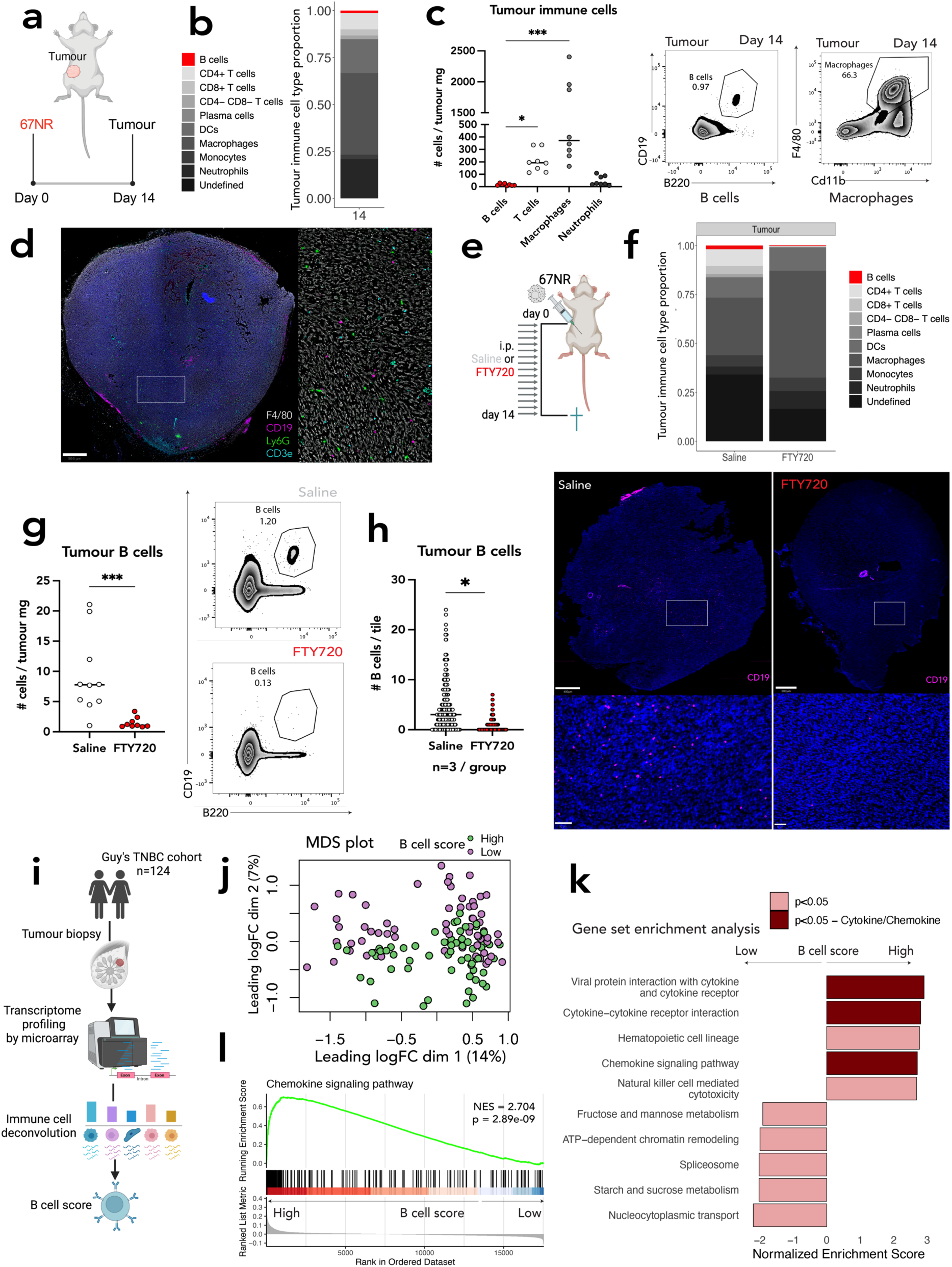
TNBC tumors exclude B cells originating from lymph nodes. **(a)** Schematic of the 67NR tumor harvesting experiment. **(b)** Immune cell composition of the 67NR tumor microenvironment shown as a bar plot, with B cells highlighted in red (day 14 post-injection). **(c)** *Left*: Quantification of tumor-infiltrating immune cell types shown as number of cells per mg of tumor (day 14 post-injection). *Right*: Representative gating strategy for identifying tumor-infiltrating B cells and macrophages (gated on CD45⁺ cells) (day 14 post-injection). **(d)** Immunofluorescence image and inset of 67NR tumors stained using a PhenoImager panel, visualizing nuclei (DAPI, blue), B cells (CD19, magenta), neutrophils (Ly6G, green), and T cells (CD3, cyan) (day 14 post-injection). Scale bar, 500 μm. **(e)** Schematic of the FTY720 treatment experiment in 67NR tumor-bearing mice. **(f)** Immune cell composition of the 67NR tumor microenvironment in mice treated with saline or FTY720, shown as a bar plot with B cells in red (day 14 post-injection). **(g)** *Left*: Quantification of tumor-infiltrating B cells (number of cells per mg of tumor) in mice treated with saline or FTY720 (day 14 post-injection). *Right:* Representative gating strategy for tumor-infiltrating B cells. **(h)** *Left*: Quantification of tumor-infiltrating B cells by immunofluorescence (number of B cells per tile) (day 14 post-injection). *Right*: Representative immunofluorescence image and inset of 67NR tumors from saline- or FTY720-treated mice, stained for B cells (CD19, magenta). Scale bars: 400 μm, 500 μm, 50 μm. **(i)** Schematic of immune cell deconvolution workflow applied to TNBC patient transcriptomes (n = 124) using microarray data. **(j)** Multidimensional scaling (MDS) plot of TNBC transcriptomes (n = 124), with each patient colored by B cell score group. **(k)** KEGG gene set enrichment analysis comparing TNBC patients by B cell score group. Top 10 enriched pathways for each group are shown; significantly enriched pathways are highlighted in pink, and cytokine/chemokine-related pathways are highlighted in rust. **(l)** Gene set enrichment plot for the KEGG pathway “Chemokine signaling pathway.” In panels (c), (g), and (j), each dot represents an individual mouse or patient [(c) *n* = 8, (h) Saline *n* = 10, FTY720 *n* = 10, (j) *n* = 124,]. In panel (h), each dot represents an individual tile and the tiles from individual mice have been combined [(m) Saline *n*=3, FTY720 *n*=3]. Bars represent the mean. Data in (b) and (c) are from the same two independent experiments. Data in (g) and (h) are from the same two independent experiments. \**P* ≤ 0.05; \*\*\**P* ≤ 0.001 [paired Friedman test with Dunn’s multiple comparison test in (c), Mann-Whitney U test in (g), nested Student’s *t* test in (h)].

To determine the source of the few B cells that do reach the tumor, we treated 67NR tumor-bearing mice with FTY720, a sphingosine-1-phosphate receptor modulator that blocks lymphocyte egress from secondary lymphoid organs^38^ (**Figure 1E; Supp Figure 1A**). FTY720 treatment markedly reduced B cell infiltration into tumors, a previously unrecognized finding that establishes the lymph node (LN) as the primary source of tumor-infiltrating B cells, alongside the expected reduction in T cells^39^, as assessed by both flow cytometry and immunofluorescence (**Figure 1F-H**; **Supp Figure 1B-E**). Thus, while B cells are capable of activation in cLNs, their recruitment to the tumor is impaired.

To explore potential mechanisms behind B cell exclusion, we analyzed transcriptomic data from 124 treatment-naïve TNBC patients^40^. Immune deconvolution^41^ permitted the interrogation of the TME in each patient, and stratification by B cell score into “Low” and “High” (**Figu**re 1I) showed that patients with higher B cell infiltration trended toward improved outcomes (**Supp Figure 1F**, Cox regression HR = 1.74, p = 0.109, CI = 0.884 – 3.422). Multidimensional scaling (MDS) indicated that B cell-high tumors clustered separately, suggesting a unique transcriptional phenotype (**Figure 1J**). Gene set enrichment analysis (GSEA) revealed that chemokine and cytokine signaling pathways were significantly enriched in these tumors (**Figure 1K-L**). These data implicate impaired chemokine signaling as a barrier to B cell trafficking in immune-cold TNBC, pointing to chemokine restoration as a rational strategy to reprogram “immune-cold” tumors.

### Chemokine supplementation fails to restore B cell migration in TNBC

Given the link between chemokine signaling and B cell infiltration in patient tumors, we next tested whether enhancing chemokine availability could overcome B cell exclusion. We first identified differentially expressed genes within the KEGG “Chemokine signaling pathway” between TNBC patients with high versus low B cell scores (**Figure 2A**). Among the top candidates were CXCL13 and CCL21, the canonical ligands for CXCR5^42,43^ and CCR7^44^, respectively; the most enriched chemokine receptors in B cell-high tumors (**Figure 2A**).

**Figure 2.**
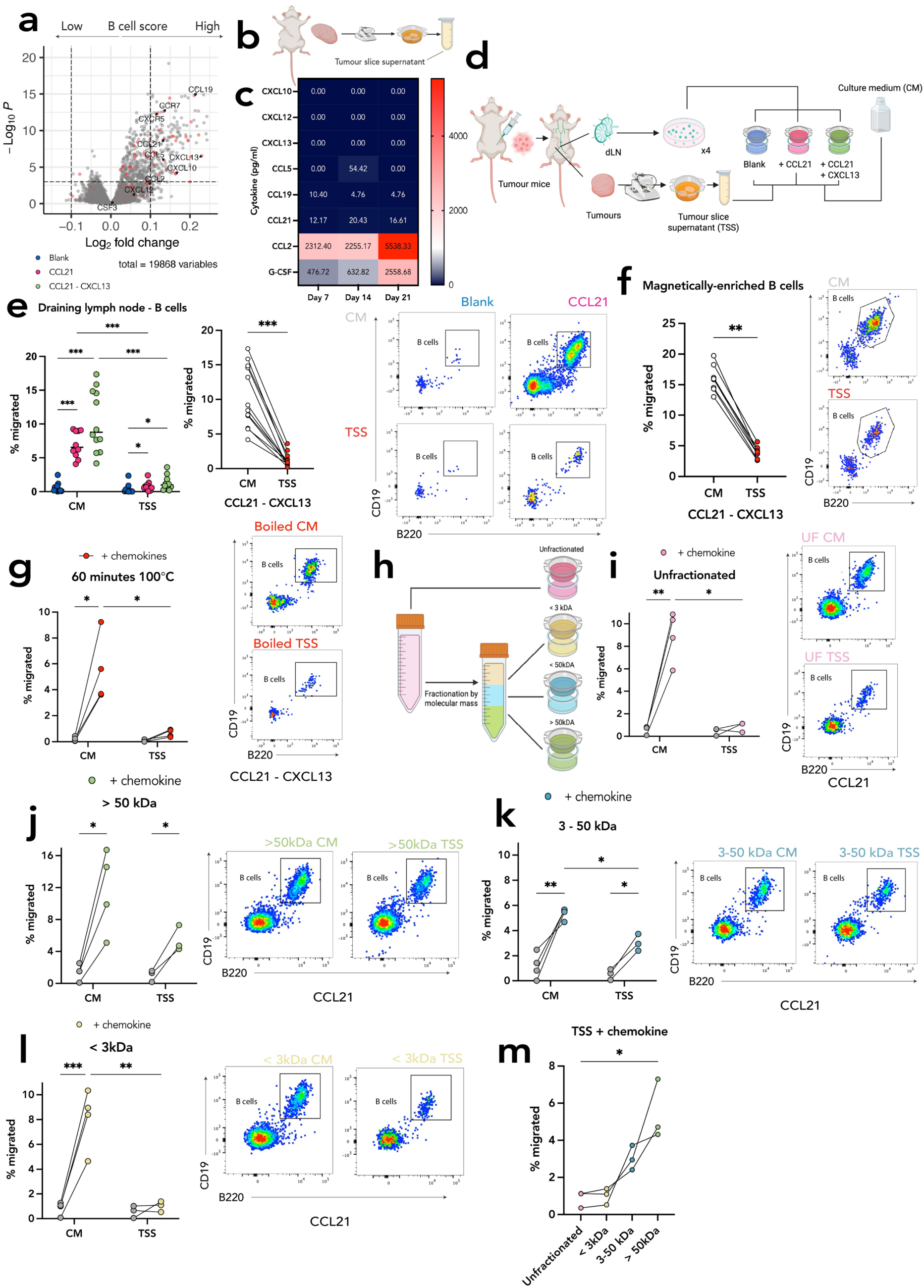
Chemokine supplementation fails to restore B cell migration in TNBC. **(a)** Volcano plot of differentially expressed genes between TNBC patients (n = 124) stratified by B cell score. Genes annotated in the KEGG “Cytokine–Cytokine Receptor Interaction” and “Chemokine Signaling” pathways are shown in red, all others in grey. **(b)** Schematic of tumor slice supernatant (TSS) generation. **(c)** Heatmap showing concentrations (pg/mL) of selected cytokines and chemokines in TSS collected at different time points post-tumor injection. **(d)** Schematic of transwell chemotaxis assay using tumor slice culture medium (CM) or TSS, with or without supplementation with CCL21 and/or CXCL13 (300 ng/mL each). **(e)** *Left*: Quantification of B cell migration (percentage of cells found in bottom transwell) towards blank or chemokine-supplemented CM and TSS. *Middle*: Paired comparison of B cell migration from lymph node suspensions towards CCL21+CXCL13-supplemented CM or TSS. *Right*: Representative gating strategy for migrated B cells under each condition. **(f)** *Left*: Paired comparison of B cell migration from magnetically enriched lymph node B cell suspensions towards chemokine-supplemented CM or TSS. *Right*: Corresponding gating strategy. **(g)** *Left:* Paired comparison of B cell migration in response to boiled CM or TSS, with or without chemokine supplementation. *Right*: Gating strategy for migrated B cells. **(h)** Schematic of TSS molecular weight fractionation using Amicon ultracentrifuge filters. **(i–l)** Paired B cell migration quantification towards blank or chemokine-supplemented CM and TSS fractions: (i) unfractionated, (j) >50 kDa, (k) 3–50 kDa, (l) <3 kDa. Right panels: Representative gating strategies for migrated B cells towards chemokine-supplemented medium. **(m)** B cell migration quantification from lymph node suspensions in response to different molecular weight fractions of chemokine-supplemented TSS. Each symbol represents an individual mouse [(e) n = 12; (f) n = 8; (g) n = 4; (i), (j), (k), (l), (m) n = 4]. Bars represent the mean. Data in (c) are from 1 independent experiment (n = 3 per time point). Data in (e) are from 3 independent experiments. Data in (f) are from 2 independent experiments. Data in (g) are from one independent experiment and data in (i), (j), (k), (l) and (m) are from the same independent experiment. \**P* ≤ 0.05; \*\**P* ≤ 0.01; \*\*\**P* ≤ 0.001. [Two-way ANOVA in left (e); paired Wilcoxon test in middle (e); paired Wilcoxon test in (f); paired two-way ANOVA in (g); paired mixed-effect analysis in (i), (j), (k), (l); paired Friedman test with Dunn’s multiple comparison correction in (m)].

To evaluate whether these chemokines are present in the TME of the TNBC mouse model, we developed an organotypic tumor slice culture system and used a custom Luminex assay to quantify chemokines in tumor slice supernatant (TSS) (**Figure 2B**). Consistent with the observed paucity of tumor-infiltrating B cells (**Figure 1B-D**), we detected very low levels of B cell-recruiting chemokines (CXCL13, CCL21), whereas cytokines linked to macrophage and neutrophil recruitment (e.g., CCL2, G-CSF) were abundant (**Figure 2C**). Notably, the chemokine profile of 67NR tumor cells cultured in isolation differed from TSS, suggesting that non-tumor components contribute to the secretome (**Supp Figure 2A**).

To test the functional relevance of these chemokine profiles, we performed transwell chemotaxis assays using whole spleen suspensions or magnetically enriched B cells. TSS induced robust neutrophil migration but failed to attract B cells (**Supp Figure 2B**). When CCL21 and/or CXCL13 were added to TSS, B cell migration remained significantly impaired compared to chemokine-supplemented control medium (**Figure 2D-E**). This inhibitory effect was independent of cell viability (**Supp Figure 2C**) and reproducible using purified B cells from naïve LNs (**Figure 3F**). These results reveal that chemokine supplementation cannot restore B cell trafficking into the TME, uncovering a previously unrecognized active barrier whereby tumor-derived factors suppress chemokine-mediated B cell migration.

**Figure 3.**
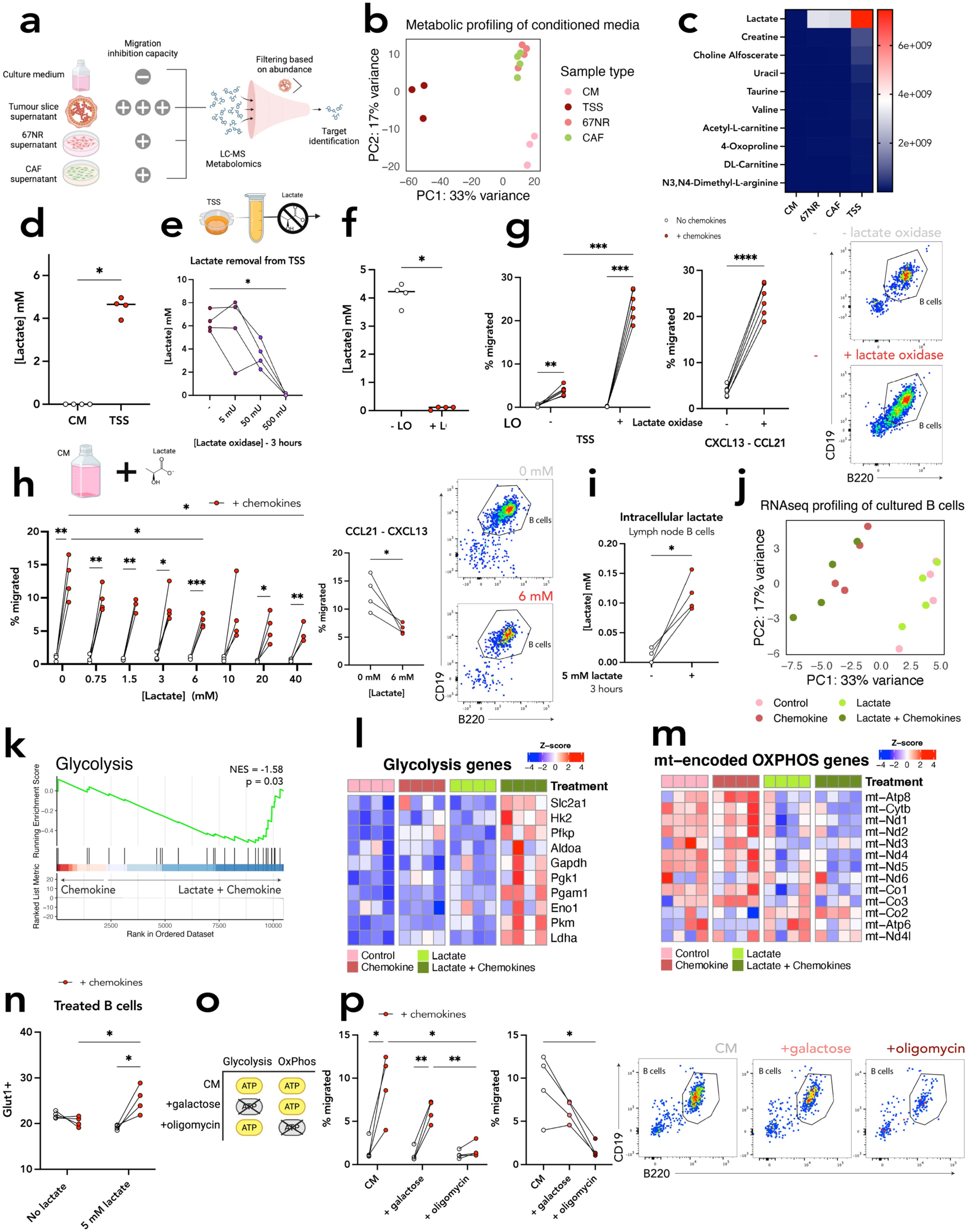
Lactate inhibits B cell migration by disrupting mitochondrial metabolism. **(a)** Schematic of the LC-MS experimental design using tumor-derived media with increasing inhibitory capacity. **(b)** Principal component analysis (PCA) of LC-MS metabolomic profiles from CM, 67NR-conditioned media, CAF-conditioned media, and TSS. **(c)** Heatmap showing the 10 most differentially abundant metabolites in TSS. **(d)** Quantification of lactate concentration in CM and TSS. **(e)** *Top*: Schematic of lactate depletion from TSS using lactate oxidase (LO). *Bottom*: Lactate quantification in TSS after 3-hour incubation with increasing concentrations of LO. **(f)** Lactate quantification in TSS before and after 3-hour incubation with 500 mU of LO. **(g)** *Left*: Quantification of B cell migration towards blank or chemokine-supplemented TSS (±LO treatment). Middle: Paired B cell migration quantification in purified B cell suspensions towards CCL21+CXCL13-supplemented TSS (±LO treatment). *Right*: Representative gating strategy for migrated B cells. **(h)** *Top*: Schematic of lactate addition to CM. Left: Quantification of B cell migration in CM with increasing lactate concentrations, with or without chemokine supplementation. *Middle*: Paired B cell migration quantification in response to CCL21+CXCL13-supplemented CM with or without 6 mM lactate. *Right*: Representative gating strategy. **(i)** Intracellular lactate levels in purified lymph node B cells after 3-hour incubation in CM with or without 5 mM lactate. **(j)** PCA of transcriptomes from cultured B cells exposed for 4 h to chemokines and/or lactate. **(k)** Gene set enrichment analysis comparing “Chemokine” vs. “Lactate + Chemokine” conditions for the Wikipathway “Glycolysis and Gluconeogenesis”. **(l)** Heatmap of scaled expression of glycolysis-related genes across B cell treatment groups. **(m)** Heatmap of scaled expression of mitochondrially encoded protein-coding genes across treatment groups **(n)** Quantification of Glut1⁺ B cells by flow cytometry after chemokine stimulation ± lactate treatment. **(o)** Schematic depicting how glucose, galactose, or oligomycin affect ATP production through glycolysis or oxidative phosphorylation (OXPHOS). **(p)** *Left*: Quantification of B cell migration in CM containing glucose (10 mM), galactose (10 mM), or oligomycin (1 μM), with or without chemokine supplementation. Middle: Paired B cell migration quantification in response to chemokine-supplemented CM containing each metabolic modulator. *Right*: Representative gating strategy. Except when otherwise stated, each symbol represents an individual mouse [(d) TSS n = 4, (e) TSS n = 4, (g) n = 6, (h) n = 4, (i) n = 4, (j) n = 4, (n) n = 4, (p) n=4]. In (b), the CM, 67NR conditioned media and CAF conditioned media are technical replicates while the TSS are biological replicates from different mice [(b) CM n = 3, CAF n = 4, 67NR n = 4, TSS n =3]. In (f), each symbol represents a technical replicate from one TSS [(d) n = 4]. Bars represent the mean. Data in (d), (e), (f), (i), (n) and (p) are from one independent experiment. Data in (g) are from two independent experiments. Data in (h) are representative of three experiments. Data in (b) and (c) are from the same independent experiment. Data in (j), (k), (l) and (m) are from the same independent experiment. \**P* ≤ 0.05; \*\**P* ≤ 0.01; \*\*\**P* ≤ 0.001. \*\*\*\**P* ≤ 0.0001. [Mann-Whitney U test in (d); paired Friedman test with Dunn’s multiple comparison correction in (e); Mann-Whitney U test in (f); paired two-way ANOVA in left (g) paired Wilcoxon test in middle (g); paired mixed effects analysis in left (h) paired Wilcoxon test in middle (h); paired Wilcoxon test in (i); paired two-way ANOVA in (n); paired two-way ANOVA in left (p), paired Friedman test with Dunn’s multiple comparison correction in middle (p)].

To investigate the nature of inhibitory factors, we first assessed heat stability. Boiling the TSS for 60 minutes failed to restore B cell migration (**Figure 2G**), indicating that the suppressive factors are heat-stable and likely not proteins. We then fractionated the TSS by molecular weight into >50 kDa, 3–50 kDa, and <3 kDa fractions using centrifugal filters (**Figure 2H**). Whereas the >50 kDa and 3–50 kDa fractions supported robust chemokine-induced B cell migration, comparable to control culture medium, the **<3** kDa fraction retained full inhibitory activity, completely blocking chemokine-driven lymphocyte migration (**Figures 2I–M**). These findings strongly suggest that a small, heat-stable, tumor-derived metabolite is responsible for suppressing B cell chemotaxis in TNBC, identifying a novel, active barrier to immune infiltration that cannot be overcome by chemokine supplementation alone.

### Lactate inhibits B cell migration by disrupting mitochondrial metabolism

To identify the tumor-derived metabolite responsible for suppressing B cell chemokine-mediated migration, we performed liquid chromatography–mass spectrometry (LC-MS) on media with varying inhibitory capacities (**Figure 3A**). These included non-inhibitory culture medium, fully inhibitory TSS, and conditioned media from 67NR tumor cells or cancer-associated fibroblasts (CAFs), which showed only mild inhibition (**Supp Figure 3A**). Principal component analysis (PCA) revealed distinct clustering, with TSS forming a unique group, separate from conditioned media and culture medium (**Figure 3B**). Among the differentially abundant metabolites, lactate was the most enriched in TSS (**Figure 3C**). This aligns with the Warburg effect, whereby tumor cells produce high levels of lactate even under aerobic conditions^45^. Measured concentrations of lactate reached ∼5 mM (**Figure 3D**), within the range reported for human tumors (2–15 mM)^46^.

To test whether lactate drives the inhibitory effect, we depleted it from <3 kDa TSS using bacterial lactate oxidase (LO), which irreversibly converts lactate to pyruvate^47^. Treatment with LO fully removed lactate (**Figure 3E-F**) and restored B cell migration in chemokine-supplemented TSS, confirming lactate as the key inhibitory component (**Figure 3G**). We next asked whether lactate alone is sufficient to impair migration. Increasing lactate concentration in culture medium suppressed chemokine-induced B cell migration in a dose-dependent manner, with 6 mM significantly reducing chemotaxis (**Figure 3H**). This effect was independent of pH, as both sodium lactate and lactic acid inhibited migration, while pH-matched culture medium had no effect (**Supp Figure 3B-E**). Importantly, none of these effects were attributable to reduced cell viability (Supp Figure 3F).

To investigate how lactate disrupts B cell migration, we first measured intracellular lactate levels and found significant uptake within 3 hours of exposure (**Figure 3I**). Transcriptomic profiling revealed that lactate had minimal impact on unstimulated B cells but altered the gene expression of chemokine-stimulated B cells (**Figure 3J**). Both chemokine-alone and chemokine plus lactate groups showed comparable upregulation of activation and signaling genes (e.g., *Cd86*, *Cd69, Jak3*, *Stat3*), confirming that chemokine-induced activation occurs even in the presence of lactate (**Supp Figure 3G**). However, lactate-treated B cells, displayed a distinct metabolic transcriptional profile upon chemokine stimulation. Glycolytic genes were strongly upregulated, while mitochondrially encoded electron transport chain genes were broadly downregulated (**Figure 3K-M**), indicating a shift from oxidative phosphorylation (OXPHOS) to aerobic glycolysis. Notably, *Ldha*, which converts pyruvate to lactate, was among the most upregulated (**Figure 3L**), suggesting that pyruvate is being diverted away from the TCA cycle, reinforcing this metabolic rerouting. This transcriptional program was accompanied by increased Glut1 surface expression (**Figure 3N**), consistent with enhanced glucose uptake^48^ and a metabolic state incompatible with mitochondrial energy production.

To determine whether this metabolic shift interferes with migration, we challenged B cells under energy-restricted conditions. Replacing glucose with galactose, which forces reliance on OXPHOS, had no effect on chemokine-induced migration, indicating that glycolysis is dispensable. In contrast, oligomycin, an OXPHOS inhibitor, completely abrogated B cell migration (**Figure 3O-P**). These findings indicate that OXPHOS, not glycolysis, is essential for B cell chemotaxis, and that lactate disrupts this requirement by inducing a glycolytic, mitochondrially-suppressed state. Supporting this, actin polymerization, a key cytoskeletal event in migration, was significantly impaired in lactate-treated B cells (**Supp Figure 3H**). Together, these data demonstrate that tumor-derived lactate directly inhibits B cell migration by rewiring their metabolic program, suppressing mitochondrial respiration required for chemokine-guided movement.

### Lactate production blockade unlocks B cell infiltration and TLS formation

Having identified tumor-derived lactate as a dominant suppressor of B cell migration, we next tested whether combining chemokine supplementation with lactate inhibition could reprogram the immune landscape of an “immune-cold” TNBC model to enable B cell infiltration and TLS formation.

To supply B cell-recruiting cues, we engineered CAFs^49^ to secrete the chemokines CXCL13 and CCL21 (**Supp Figure 4A-B**). ELISA confirmed robust chemokine production both *in vitro* and *in vivo*, with detectable levels in cell culture supernatants, tumor homogenates and interstitial fluid following co-injection of CAFs with 67NR TNBC cells (**Supp Figure 4C-F**). However, chemokine delivery alone failed to enhance B cell infiltration (**Supp Figure 4G-H**), indicating that the suppressive effect of lactate persists *in vivo* and may override chemokine-driven recruitment.

**Figure 4.**
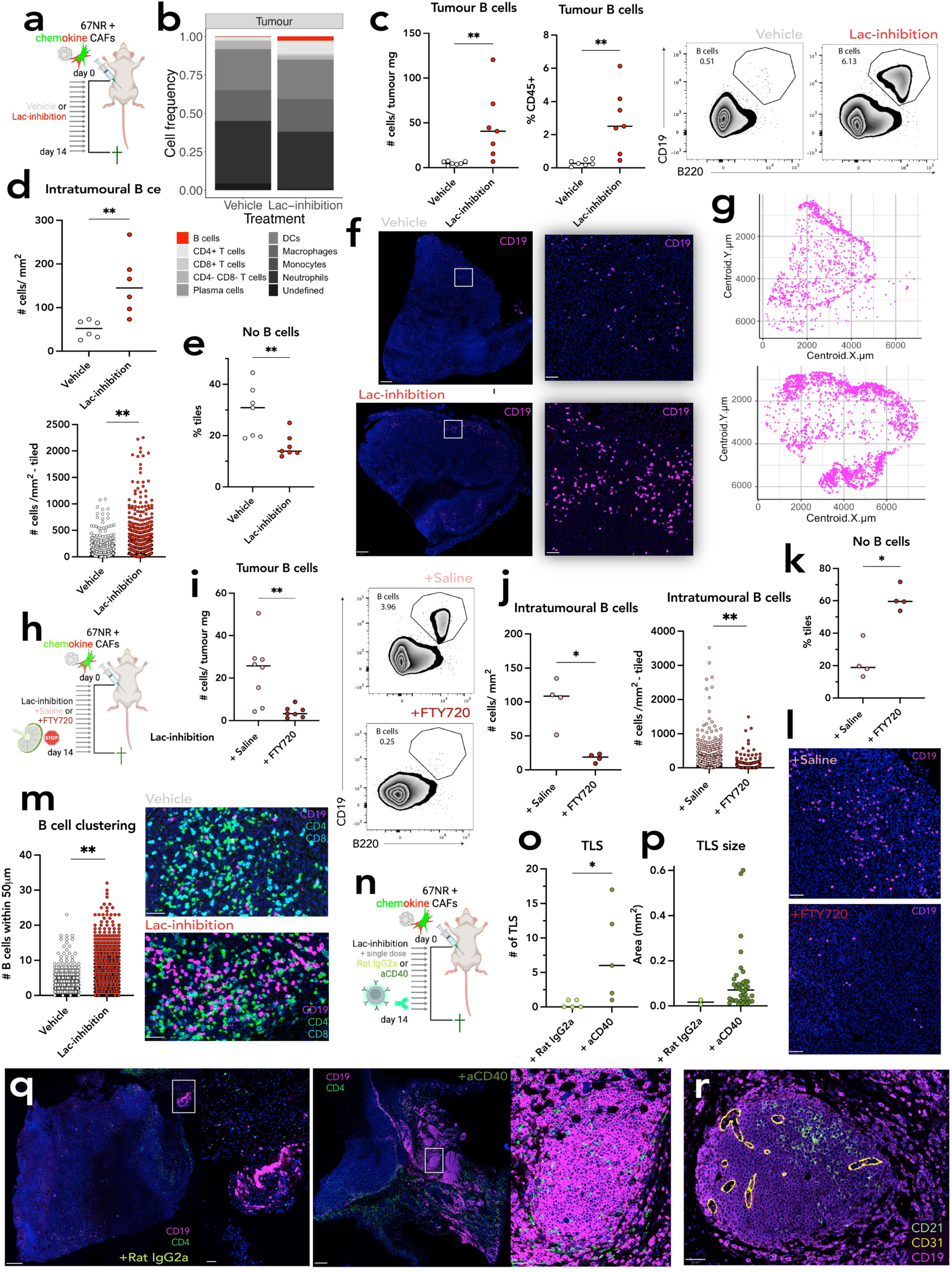
Lactate production blockade unlocks B cell infiltration and TLS formation. **(a)** Schematic of the lactate production inhibition experiment in mice bearing 67NR + chemokine CAF tumors. **(b)** Immune cell composition of the 67NR+chemokine CAF tumor microenvironment, with or without lactate inhibition, shown as a bar plot. B cells are highlighted in red (day 14 post-injection). **(c)** *Left and middle*: Quantification of tumor-infiltrating B cells by flow cytometry, shown as number per mg tumor *(left)* and percentage of CD45⁺ cells (*middle*), in the presence or absence of lactate inhibition. *Right*: Representative gating strategy for B cells (gated on CD45⁺ cells) (day 14 post-injection). **(d)** *Top:* Quantification of B cell infiltration by immunofluorescence, shown as number of B cells per mm². *Bottom:* Tiled B cell density analysis (each dot = one tile). **(e)** Tumor B cell desertification shown as % of tiles with 0 B cells, with or without lactate inhibition. **(f)** Representative immunofluorescence images of 67NR+chemokine CAF tumors with or without lactate inhibition, stained for CD19⁺ B cells (magenta). Left: Whole-tumor view. Right: Insets showing magnified tumor regions. Scale bars: 500 μm, 50 μm. **(g)** Digital tissue maps of segmented B cells in tumors represented in (f) generated using RStudio from QuPath measurements. **(h)** Schematic of the FTY720 treatment experiment in 67NR+chemokine CAF tumor-bearing mice under lactate inhibition. **(i)** *Left:* Quantification of tumor-infiltrating B cells by flow cytometry (cells per mg tumor) in mice treated with lactate inhibition plus saline or FTY720 (day 14 post-injection). *Right:* Representative gating strategy. **(j)** Quantification of B cells by immunofluorescence, shown as number of B cells per mm² (*left*) and tiled B cell density (*right*). **(k)** Tumor B cell desertification shown as % of tiles with 0 B cells, with or without FTY720 treatment under lactate inhibition. **(l)** Representative immunofluorescence images of CD19⁺ B cells (magenta) in tumor areas from saline- or FTY720-treated mice under lactate inhibition. Scale bars: 50 *μ* m. **(m)** *Left*: B cell clustering quantified by immunofluorescence (number of neighboring B cells within 50 μm per B cell), with or without lactate inhibition. *Right*: Representative immunofluorescence images showing CD19⁺ B cells (magenta), CD4⁺ T cells (green), and CD8⁺ T cells (cyan). Scale bars: 100 μm. **(n)** Schematic of the single-dose agonistic anti-CD40 (aCD40) treatment experiment in mice bearing 67NR+chemokine CAF tumors under lactate inhibition. **(o)** Quantification of TLS numbers in mice treated with isotype control (Rat IgG2a) or aCD40. **(p)** Quantification of individual TLS area in mice treated with Rat IgG2a or aCD40. **(q)** Immunofluorescence images showing TLS in 67NR+chemokine CAF tumors under lactate inhibition following Rat IgG2a or aCD40 treatment, stained for CD19⁺ B cells (magenta) and CD4⁺ T cells (green). Insets highlight TLS structures. Scale bars: 400 μm, 50 μm. **(r)** Immunofluorescence image showing TLS structure in an aCD40-treated 67NR+chemokine CAF tumor under lactate inhibition stained for B cells (CD19, magenta), follicular dendritic cells (FDCs) (CD21, green) and blood vessels (CD31, yellow). Scale bars: 50 μm. Except when otherwise stated, each symbol represents an individual mouse [(c) Vehicle n = 7, Lac-inhibition n = 7; (d) Vehicle n = 6, Lac-inhibition n = 6; (e) Vehicle n = 6, Lac-inhibition n = 6; (i) +Saline n = 8, +FTY720 n = 8; (j) +Saline n = 4, +FTY720 n = 4; (k) +Saline n = 4, +FTY720 n = 4; (o) +Rat IgG2a n = 5; +aCD40 n = 5]. In bottom (d) and right (j), each dot represents an individual tile and the tiles from individual mice have been combined [top (d) Vehicle n = 7, Lac-inhibition n = 7; right (j) +Saline n = 4, +FTY720 n=4]. In (m), each dot represents an individual B cell and the B cells from individual mice have been combined [(m) Vehicle n = 6, Lac-inhibition n = 6]. In (p), each dot represents an individual TLS and TLS from individual mice have been combined [(p) +Rat IgG2a n=2, +aCD40 n=5]. Bars represent the mean. Data in (b), (c), (d), (e) and (m) are from the same two independent experiments. Data in (i) are from two independent experiments, including one experiment used for (j) and (k). Data in (o) and (p) are from the same independent experiment. \**P* ≤ 0.05; \*\**P* ≤ 0.01. [Mann-Whitney U test in (c), top (d), (e), (i), left (j), (k) and (o); nested Student’s t test in bottom (d), right (j), (m) and (p)].

To test this, we pharmacologically inhibited lactate production using the lactate dehydrogenase (LDH) inhibitor oxamate in mice bearing 67NR plus Chemokine-CAF tumors (**Figure 4A**). Lactate blockade significantly enhanced B cell infiltration, as shown by flow cytometry (**Figure 4B-C**) and immunofluorescence (**Figure 4D-G**). B cells were no longer confined to the tumor periphery but were broadly distributed throughout the tumor bed (**Figure 4F-G)**, reaching up to 6% of CD45⁺ cells (**Figure 4B-C**). These findings demonstrate that combining chemokine supplementation with lactate inhibition overcomes B cell exclusion and promotes a more “inflamed” immune landscape.

To confirm the origin of infiltrating B cells, we repeated the experiment with FTY720 to block lymphocyte egress from LNs (**Figure 4H**). FTY720 treatment abrogated circulating B cells (**Supp Figure 4I**) and significantly reduced B cell infiltration into tumors, as measured by flow cytometry and immunofluorescence (**Figure 4I-L**), confirming that B cells were actively recruited from lymphoid tissues under these conditions. Spatial analysis further revealed a shift back to the “immune-ignored” tumor phenotype in FTY720 treatment condition, with nearly 60% of tumor regions completely lacking B cells (**Figure 4K**). These results confirm that lactate inhibition promotes B cell chemokine-mediated infiltration via enhanced trafficking from LNs.

Notably, tumor-infiltrating B cells in lactate-inhibited tumors frequently clustered with other B cells, although clustering with CD4⁺ or CD8⁺ T cells remained unchanged (**Figure 4M**; **Supp Figure 4J**). B-cell-to-B-cell clustering was significantly increased, raising the possibility of TLS formation, as such structures typically arise from dense aggregates of B cells^50^. To test this, we combined lactate inhibition and chemokine supplementation with a single dose of agonistic anti-CD40 antibody (aCD40), which mimics T cell help and broadly activates B cells^51^ (**Figure 4N**). While aCD40 only mildly increased the total number of tumor-infiltrating B cells (**Supp Figure 4K-L**), it significantly increased TLS frequency (isotype control 2/5 vs. aCD40: 5/5) (**Supp Figure 4M**), number (**Figure 4O**), and size (**Figure 4P**). The largest TLS exceeded 0.5 mm² and contained over 4000 cells (**Figure 4P**; **Supp Figure 4N**). In aCD40-treated mice, immunofluorescence revealed large, well-organized peritumoral B cell aggregates with sparse CD4⁺ T cell infiltration (**Figure 4Q**), and multiplex staining showed CD21⁺ networks and CD31⁺ vessels in some TLS, consistent with the presence of follicular dendritic cells and vascularization (**Figure 4R**).

Together, these results demonstrate that chemokine delivery combined with lactate inhibition is sufficient to induce B cell infiltration and TLS formation, and that B cell stimulation via CD40 induces TLS maturation by further amplifying this response in an immune-excluded breast cancer model. This approach offers a tractable strategy to reprogram immune-cold tumors into TLS-rich, immunologically active microenvironments.

### Lactate antagonizes B cell infiltration and TLS Formation across human cancers

To investigate whether tumor-derived lactate limits B cell infiltration in patients, we used a glycolysis-related gene expression signature “Aerobic Glycolysis (AG)” (WP4629^52^) as a proxy for lactate production. Across multiple cancer cohorts, transcriptomic profiles were stratified by glycolytic activity (“High AG” vs. “Low AG”) and further grouped by *CXCL13* expression to capture chemokine availability. This allowed us to examine how metabolic and chemokine signals interact to shape immune infiltration in the TME (**Figure 5A**).

**Figure 5.**
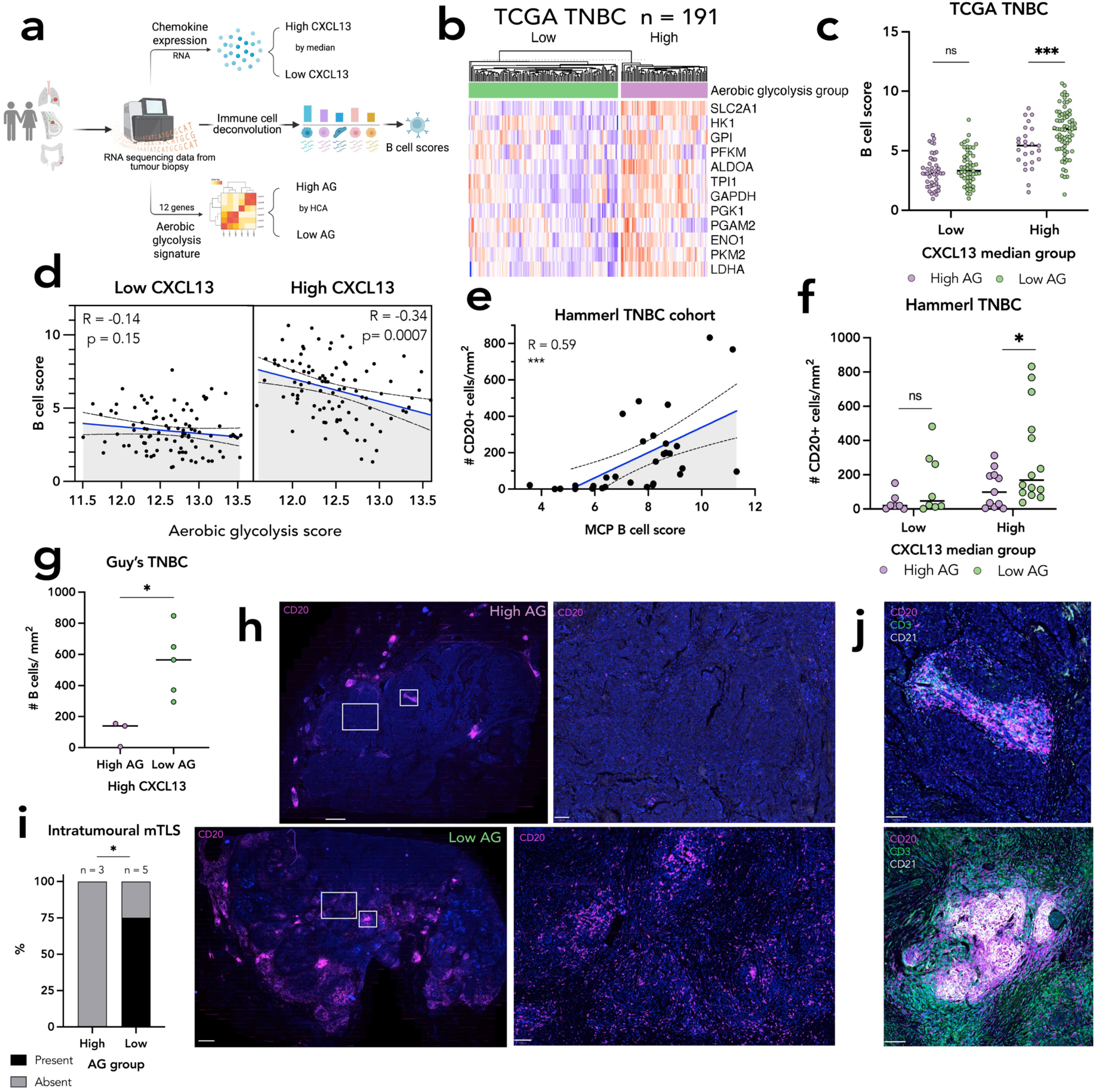
Lactate antagonizes B cell infiltration and TLS Formation across human cancers. **(a)** Schematic of analytical workflow for TNBC patient cohorts, including stratification by median CXCL13 expression, hierarchical clustering based on an Aerobic Glycolysis (AG) gene signature, and immune cell deconvolution. **(b)** Heatmap showing scaled expression of 12 AG-related genes in TCGA TNBC patients (n = 191), grouped by hierarchical clustering. **(c)** B cell scores (MCP-Counter analysis) across four CXCL13/AG groups in the TCGA TNBC cohort. **(d)** Correlation between AG score and B cell score (MCP-Counter) in TCGA TNBC patients, shown separately for CXCL13-high and CXCL13-low groups. **(e)** Correlation between B cell scores (MCP-Counter) and histological CD20⁺ cell density (cells/mm²) in the Hammerl TNBC cohort (n = 35). **(f)** Histological CD20⁺ B cell density across the four CXCL13/AG groups in the Hammerl TNBC cohort. **(g)** Quantification of tumor-infiltrating B cells (CD20⁺ cells/mm²) by immunofluorescence in CXCL13-high TNBC patients from Guy’s cohort, stratified by AG cluster. **(h)** Representative immunofluorescence images (with insets) of TNBC tumors from Guy’s cohort stained for B cells (CD20, magenta), illustrating examples from each AG cluster. **(i)** Contingency plot showing the frequency of intratumoral mature tertiary lymphoid structures (mTLS), defined by the presence of CD21⁺ follicular dendritic cells, in CXCL13-high TNBC patients from Guy’s cohort across AG clusters. **(j)** Representative immunofluorescence images of TLS stained for B cells (CD20, magenta), T cells (CD3, green), and follicular dendritic cells (CD21, white). Each symbol represents an individual patient [(c) TCGA TNBC: Low CXCL13 High AG n = 45, Low CXCL13 Low AG n = 51, High CXCL13 High AG n = 25, High CXCL13 Low AG n = 70; (d) TCGA TNBC: Low CXCL13 n = 96, High CXCL13 n = 95; (e) Hammerl TNBC: n = 31; (f) Hammerl TNBC: Low CXCL13 High AG n = 7, Low CXCL13 Low AG n = 9, High CXCL13 High AG n = 12, High CXCL13 Low AG n = 14; (g) Guy’s TNBC: High AG n = 3, Low AG n = 5]. Bars represent the mean. Data in (c) and (g) are from one independent experiment. Data in (g) and (i) are from the same independent experiment. *P ≤ 0.05; **P ≤ 0.01; ***P ≤ 0.001. [Two-way ANOVA in (c) and (f), only results within CXCL13 groups are shown; Pearson’s correlation test in (d) and (e); Mann-Whitney U test in (g); Chi-square test in (i)].

In the TCGA TNBC cohort^53^ (n=191), B cell gene expression scores were significantly higher in CXCL13-high patients with low AG, compared to those with high AG (**Figure 5B-C**). This result was consistent across immune deconvolution algorithms (**Supp Figure 5A**) and was absent in CXCL13-low patients. Supporting a metabolic-chemokine interaction, AG signature and B cell scores were significantly and inversely correlated only in CXCL13-high tumors (**Figure 5D**). These findings were validated in the SCANB TNBC cohort^54^ (n=202), and in the larger ISPY-2 pan-breast cancer dataset^55^ (n=984), where B and T cell scores were again highest in CXCL13-high / Low AG tumors, with an inverse correlation between AG and B cell score specific to CXCL13-high patients (**Supp Figures 5B-G**).

To test whether this metabolic–chemokine interaction extends beyond breast cancer, we applied the same analysis to non-small cell lung cancer^56^ (NSCLC, n=924) and colon cancer^57^ (n=585) cohorts. In both, B cell infiltration was consistently highest in CXCL13-high / Low AG tumors, and AG scores were inversely correlated with B cell abundance. Notably, this relationship was again specific to CXCL13-high tumors (**Supp Figures 5H-M**).

To validate these transcriptomic findings spatially, we analyzed a TNBC cohort (n=42) with matched multiplex immunofluorescence data^10^. B cell scores strongly correlated with CD20⁺ B cell density (**Figure 5E**), and CXCL13-high / Low AG tumors exhibited significantly greater B cell infiltration by image analysis (**Figure 5F**). This pattern was confirmed in an independent TNBC cohort^40^ (Guy’s, n=8), where CXCL13-high tumors with low AG showed significantly higher intratumoral B cell density (**Figure 5G**). Spatial mapping revealed distinct infiltration patterns: B cells in High AG tumors were confined to the periphery, whereas in low AG tumors, they were distributed throughout the tumor bed, consistent with “excluded” versus “inflamed” phenotypes (**Figure 5H**).

To explore spatial dynamics further, we generated nanoString/Bruker GeoMx spatial transcriptomics from four TNBC patients (Tianjin). Within tumors, immune-rich regions of interest (ROIs) were characterized by high CXCL13 expression and low AG scores (**Supp Figure 5N-O**). ROIs were classified as immune desert, infiltrated, aggregate, or TLS based on CD20 and CD3 staining (**Supp Figure 5P**). CXCL13 expression progressively increased, and AG scores decreased along this continuum (Desert → Infiltrated → Aggregate → TLS), supporting a model in which local chemokine levels and metabolic activity jointly determine B cell positioning and organization (**Supp Figure 5Q**).

Finally, we assessed TLS formation across AG strata in CXCL13-high TNBC patients (Guy’s, n=8). Strikingly, mature intratumoral TLS (mTLS) were found only in low AG tumors (**Figure 5I-J**). This suggests that elevated tumor glycolysis not only impairs B cell infiltration but also limits their spatial organization and maturation into TLS, mirroring the lactate-driven exclusion phenotype observed in the TNBC mouse models.

## Discussion

This study identifies the cLN as the primary source for tumors-infiltrating B cells and uncovers a previously unrecognized mechanism by which tumors-derived lactate blocks this recruitment route by actively suppressing chemokine-driven migration. While the immunomodulatory properties of lactate are increasingly appreciated^58–60^, and its drainage to cLNs has been reported^61^, our work provides the first demonstration that lactate directly impairs cLN B cell chemotaxis by rewiring their metabolic programming, thereby blocking their infiltration into tumors and preventing TLS formation.

Although the Warburg effect has long been implicated in tumor proliferation and immune evasion^62^, its specific impact on B cell trafficking had not been studied. We show that lactate, abundantly secreted by glycolytic tumors, directly inhibits chemokine-guided B cell migration by disrupting mitochondrial OXPHOS, a metabolic pathway we identify as essential for B cell chemotaxis. This stands in contrast to CD4⁺ T cells, whereby lactate also suppresses chemokine-mediated migration via metabolic reprogramming, but through a distinct mechanism, namely by inhibiting glycolysis which is required for CD4^+^ T cell motility^63^.

Mechanistically, lactate uptake reprogrammed chemokine-stimulated B cells by inducing a glycolytic transcriptional profile while suppressing mitochondrial respiratory programs, including electron transport chain components, resembling the metabolic phenotype of aged, immunosenescent B cells^64^. This metabolic shift disrupted actin polymerization and impaired migration. The critical requirement for OXPHOS in B cell chemotaxis was confirmed by the inability of oligomycin-treated B cells to respond to chemotactic cues. These findings support emerging evidence that mitochondrial fitness, including transcription and translation of mitochondrial DNA, is essential for B cell motility and effective germinal center responses^65^. Whether lactate-mediated inhibition of B cell migration extends to other chemoattractant receptor-ligand systems^66^ remains unknown.

Our study uncovers a novel mechanism by which cancer cells can blunt chemokine-orchestrated organization of the TME^67^. These findings offer an explanation for the failure of chemokine supplementation to enhance B cell infiltration in preclinical models^68–71^, despite strong clinical associations between CXCL13 and TLS presence^72–74^. Our work suggests this is due to the overriding effect of lactate, which blocks chemokine responsiveness at the cellular level. Importantly, combining chemokine delivery with pharmacologic inhibition of lactate production re-enabled B cell trafficking and permitted TLS induction. Site-specific variation in TLS formation and composition across tumor models has long posed a challenge to TLS research^75^. Our data offer a unifying explanation: subcutaneous and mammary fat pad tumors exhibit higher lactate levels due to hypoxia and poor perfusion^76^, both of which promote lactate accumulation and suppress B cell infiltration, which may function as a bottleneck to TLS formation. This may explain the rarity of tumor-associated TLS in breast cancer models, compared to intraperitoneal^75,77^ or orthotopic sites like lung^78,79^ and brain^80^, which support B cell-rich aggregates more readily.

We further demonstrate that TLS formation is not solely contingent on B cell presence. While lactate blockade and chemokine delivery permitted B cell clustering, the addition of a single dose of agonistic anti-CD40 antibody markedly increased TLS number and size. This suggests that recruitment must be paired with immune activation to achieve full TLS maturation. This aligns with other studies showing that TLS formation in B cell-permissive environments depends on additional immunostimulatory factors such as increased antigenicity^81^ or relief from immunosuppression^82^. This reinforces the concept that TLS formation is a multi-step process requiring both cellular entry and appropriate local cues^83^.

Translating these findings to human tumors, we found that the combination of high CXCL13 expression and low glycolytic activity was strongly associated with B cell infiltration across breast, lung, and colon cancers and intratumoral mature TLS presence in TNBC. These associations, confirmed in multiple independent cancer datasets and by spatial transcriptomics assessment of TNBC, suggest that B cell infiltration is constrained by a metabolic-chemokine axis conserved across tumor types.

Functionally, TLS are increasingly recognized as intratumoral immune hubs capable of supporting antigen presentation, lymphocyte activation, and antibody production. Promoting their formation may thus represent a strategy to convert immunologically “cold” tumors into “hot,” therapy-responsive environments. Our data suggest that successful TLS induction in excluded tumors requires a coordinated approach that: (1) removes metabolic barriers (e.g., through lactate blockade), (2) restores chemokine-mediated recruitment, and (3) provides inflammatory stimulation (e.g., CD40 agonism). This tripartite strategy could offer a rational blueprint for designing therapies to overcome immune exclusion.

Given that high serum LDH is associated with poor ICI response^84^, while CXCL13 and TLS are linked to improved outcomes^73,74^, combining lactate inhibition, chemokine delivery, and immune activation may synergistically improve immunotherapy efficacy in TNBC and other tumors characterized by B cell exclusion.

## Materials & Methods

### Patient Sample Collection

Tumor transcriptome profiling via microarray was obtained from 124 treatment-naïve TNBC (ER-negative, HER2-negative by IHC) patients (Guy’s cohort) treated between 1984 and 2002 at Guy’s Hospital, London, UK.

Treatment-naïve TNBC breast tumor tissue for immunofluorescence and spatial transcriptomics was provided by King’s Health Partners (KHP) Cancer Biobank, and Tianjin Medical University Cancer Institute. All tissue procurement was approved by the following research ethics committees (KHP Cancer Biobank REC: reference 18/EE/0025; Tianjin Medical University Cancer Institute and Hospital: reference Ek2020021). Formalin-fixed paraffin-embedded (FFPE) tumor blocks from 8 TNBC patients (Guy’s cohort) were included for immunofluorescence analysis and from 4 TNBC patients (Tianjin cohort) were included for spatial transcriptomics. Tissue sections (5 μm) were prepared by the KHP Cancer Biobank and at the Tianjin Medical University Cancer Institute.

### Mice

BALB/c female mice aged 7–13 weeks were used in this study. All mice were bred and maintained under specific-pathogen-free conditions at the Francis Crick Institute Biological Research Facility, in accordance with UK Home Office guidelines and approved by the Francis Crick Institute Ethical Review Panel.

### Antibodies and reagents

Anti-mouse CD4 (Clone EPR19514; Abcam; Cat#ab183685; RRID: AB_2686917); Anti-mouse CD8 (Clone EPR21769; Abcam; Cat#ab217344, RRID: AB_2890649);

Anti-mouse CD19 (Clone EPR23174-145; Abcam; Cat#ab245235, RRID: AB_2895109); Anti-mouse CD3 (Clone CD3-12; Abcam; ab11089, RRID: AB_2889189); Alexa Fluor® 647 anti-human CD20 (Clone EP459Y; Abcam; Cat#ab198943, RRID: AB_2905499); Anti-human CD21 (Clone EP30393; Abcam; Cat#ab75985, RRID: AB_1523292); Bond anti-rabbit Polymer RTU (Leica Biosystems; Cat#RE7260-CE); Alexa Fluor® 647 Affinipure® Donkey Anti-Rabbit IgG (H+L) (Polyclonal; Jackson ImmunoResearch; Cat#711-605-152; RRID: AB_2492288); Alexa Fluor® 488 Affinipure® Goat Anti-Rat IgG (H+L) (Polyclonal; Jackson ImmunoResearch; Cat# 112-545-003; RRID: AB_2338351); Alexa Fluor® 488 Affinipure® Donkey Anti-Rabbit IgG (H+L) (Polyclonal; Jackson ImmunoResearch; Cat#711-545-152; RRID: AB_2313584); Anti-mouse LY6G (Clone E6Z1T; Cell Signaling Technology; Cat#87048S; RRID:AB_2909808); Anti-mouse F4/80 (Clone EPR26545-166; Abcam; Cat# ab300421; RRID:AB_2936298); Anti-mouse aSMA (Clone 1A4; Agilent; Cat#M0851; RRID: AB_2223500); Anti-mouse CD3 (Clone EPR4517; Abcam; Cat#ab134096); Anti-mouse CD45 (Clone EPR23174-145; Abcam; Cat#ab245235; RRID:AB_2895109); APC anti-mouse CD19 (Clone 6D5; BioLegend; Cat#115511; RRID:AB_313647); BV605 anti-mouse CD19 (Clone 6D5; BioLegend; Cat#115540; RRID:AB_2563067); PE-CF594 anti-mouse CD19 (Clone 1D3; BD Biosciences; Cat#562291, RRID:AB_11154223); BV711 anti-mouse B220 (Clone RA3-6B2; BioLegend; Cat#103255, RRID:AB_2563491); BUV395 anti-mouse B220 (Clone RA3-6B2; BD Biosciences; Cat#563793, RRID:AB_2738427); BUV737 anti-mouse B220 (Clone RA3-6B2; BD Biosciences; Cat#612838, RRID:AB_2870160); PerCP-Cy5.5 anti-mouse B220 (Clone RA3-6B2; BioLegend; Cat#103235; RRID:AB_893356); PE-Cy7 B220 (Clone RA3-6B2; BioLegend; Cat#103221; RRID:AB_313004); BV650 anti-mouse B220 (Clone RA3-6B2; BD Biosciences; Cat#563893; RRID:AB_2738471); BV605 anti-mouse CD45 (Clone 30-F11; BioLegend; Cat#103139; RRID:AB_2562341); PE anti-mouse CD11c (Clone N418; Thermo Fisher Scientific; Cat#25-0114-81; RRID:AB_469589); PE-Cy7 anti-mouse CD3e (Clone 145-2C11; BioLegend; Cat#100320; RRID:AB_312685); Alexa Fluor 488 anti-mouse CD3e (Clone 145-2C11; BioLegend; Cat#100321; RRID:AB_389300); Biotin anti-mouse CD3e (Clone 145-2C11; BioLegend; Cat#100303; RRID:AB_312668); BUV395 anti-mouse CD3e (Clone 145-2C11; BD Biosciences; Cat#563565; RRID:AB_2738278); PerCP-Cy5.5 anti-mouse Gr-1 (Clone RB6-8C5; BioLegend; Cat#108427; RRID:AB_893561); Alexa Fluor 700 anti-mouse Gr-1 (Clone RB6-8C5; BioLegend; Cat#108421; RRID:AB_493728); BV421 anti-mouse CD11b (Clone M1/70; BioLegend; Cat#101235; RRID:AB_10897942); PE anti-mouse CD11b (Clone M1/70; BioLegend; Cat#101207; RRID:AB_312790); BUV395 anti-mouse CD8a (Clone 53-6.7; BD Biosciences; Cat#563786; RRID:AB_2732919); BV650 anti-mouse CD8a (Clone 53-6.7; BioLegend; Cat#100741; RRID:AB_11124344); BV785 anti-mouse F4/80 (Clone BM8; BioLegend; Cat#123141; AB_2563667); APC anti-mouse CD138 (Clone 281-2; BioLegend; Cat#142506; RRID:AB_10962911); BUV737 anti-mouse CD4 (Clone GK1.5; BD Biosciences; Cat#612761; RRID:AB_2870092); Alexa Fluor 700 anti-mouse CD4 (Clone GK1.5; BioLegend; Cat#100429; RRID:AB_493698); Alexa Fluor 488 anti-mouse Glut1 (Clone EPR3915; Abcam; Cat#ab195359; RRID:AB_2714026); BV510 anti-mouse Ly6G (Clone 1A8; BioLegend; Cat#127633; RRID:AB_2562937); Fixable Viability Dye (Thermo Fisher Scientific); AffiniPure F(ab’)2 Fragment Goat Anti-Mouse IgG + IgM (H+L) (Jackson ImmunoResearch; Cat#115-006-068; RRID: AB_2338471); RNeasy Mini Kit (Qiagen); Recombinant Mouse CXCL13 (Peprotech); Recombinant Mouse CCL21 (Peprotech), Recombinant Mouse BAFF (Bio-techne); Alexa Fluor 647-labelled Phalloidin (Thermo Fisher Scientific)

### Cells

67NR and 4T1.2 cell lines (kindly provided by Tony Ng, King’s College London) were cultured in DMEM (Gibco) or RPMI 1640 (Gibco), respectively, supplemented with 10% fetal bovine serum (FBS) (Gibco) and 1% penicillin–streptomycin (Gibco). Cancer-associated fibroblasts (CAFs), generously provided by Valeria Poli (University of Turin), were maintained in DMEM (Gibco) with 10% FBS (Gibco) and 1% penicillin–streptomycin (Gibco).

### Virus production

Lentiviral particles were produced by co-transfecting 2 × 10⁷ HEK293T cells per 15-cm dish with 8.3 μg of transfer plasmid, 9.6 μg pLP1, 9.0 μg pLP2, and 6.4 μg VSV-G using 100 μL Lipofectamine 2000 (Thermo Fisher), according to the manufacturer’s instructions. Viral supernatants were collected at 48-and 72-hours post-transfection, pooled, and concentrated with 10% PEG8000 (Sigma)/88 mM NaCl (Sigma). Aliquots were stored at −80 °C. Viral titers were determined by droplet digital PCR (ddPCR) as described by Wang et al^85^.

AAV vectors were produced in HEK293T cells by triple transfection with pHelper, pAAV2/1 (a gift from James M. Wilson, Addgene #112862; RRID:Addgene_112862), and transfer plasmid in a 1:1.3:1 molar ratio using PEI-MAX. Viral supernatants were collected at 72 and 120 hours post-transfection, precipitated with 8% PEG8000 (Sigma)/82 mM NaCl (Sigma), and centrifuged. Pellets were treated with Denarase (c-Lecta Gmbh), extracted with chloroform (Sigma), and purified by iodixanol gradient (Serumwerk) ultracentrifugation. The 40% fraction was filtered and concentrated using a 100-kDa Amicon filter (Sigma). Titers were quantified by SYBR Green qPCR as described in Aurnhammer et al^86^.

SB100X in pCAG-globin-pA (a gift from Mark Groudine, Addgene #127909; RRID: Addgene_127909) was used to express the transposase gene.

### Generation of chemokine-secreting CAFs

To generate chemokine-secreting CAFs (Chemokine-CAFs), wild-type (WT) CAFs^49^ underwent two rounds of transduction. First, WT CAFs were transduced with a lentivirus encoding the CCL21-GFP construct. These CCL21-CAFs were then transduced with two adeno-associated viruses (AAVs): one encoding the Sleeping Beauty transposase and the other carrying the CXCL13-mCherry construct flanked by Sleeping Beauty transposon repeats.

For transduction, 96-well plates were coated with Retronectin (50 μg/ml) (Takara), incubated at room temperature for 2 hours, and rinsed once with PBS (Gibco). For lentiviral transduction, 10 MOI of lentivirus was added per well, followed by 1×10⁵ fibroblasts in 100 μl medium. For AAV transduction, 2 μl of neat viral suspension was added to 1×10⁵ fibroblasts. Plates were briefly centrifuged and incubated for 48 hours in DMEM (Gibco) with 10% FBS (no antibiotics) (Gibco). Selection of successfully transduced cells was performed using Geneticin (Gibco) for the lentiviral construct and puromycin (Sigma) for the AAV construct. After two weeks in selective media, CAFs were sorted by FACS on a FACSAria (BD Biosciences). CCL21-CAFs were gated as GFP^hi^, and CXCL13-CCL21 CAFs as GFP^hi^/mCherry^hi^. Fluorescence expression was monitored throughout using the EVOS Cell Imaging System (Invitrogen).

### Tumor models

For orthotopic breast cancer models, mice were anesthetized with isoflurane, and 4×10⁵ 67NR or 4T1.2 cells were injected in Matrigel (Corning) into the fourth mammary fat pad. In some experiments, 4×10⁵ 67NR cells were mixed with 4×10⁵ CAFs prior to injection. Mice were monitored for tumor growth and sacrificed at designated time points.

### Treatments

For lymph node egress blockade, mice received daily intraperitoneal (i.p.) injections of FTY720 (250 μg/mouse) (Sigma) or saline, beginning on the same day they were injected with 67NR or 67NR+CAFs.

For inhibition of lactate production, mice received daily i.p. injections of either 750 mg/kg oxamate (Lac-inhibition group) (Santa Cruz Biotechnologies) or saline (vehicle control group), beginning on the same day they were injected with 67NR+CAFs.

For antibody treatments, mice received a single i.p. dose of 200 μg anti-CD40 (Ultra-LEAF Purified, clone 3/23, BioLegend) or isotype control (Ultra-LEAF Purified Rat IgG2a, clone RTK2758, BioLegend).

### Cell preparation

To confirm FTY720 activity, peripheral blood was collected from the saphenous vein at designated time points in tumor-bearing mice treated with saline or FTY720 (Sigma). A volume of 10 μL blood per mouse was treated with ACK buffer (Gibco) for red blood cell lysis.

For tumor immunophenotyping, tumors were finely minced using a straight-edge razor and digested using the Tumor Dissociation Kit, Mouse (Miltenyi) following the manufacturer’s protocol. Tumor cell suspensions were filtered through a 70 μm filter, washed with FACS buffer (PBS, 2% FBS (Gibco), 2 mM EDTA (Gibco)), counted, and CD45⁺ immune cells were enriched using CD45 MicroBeads (Miltenyi), per the manufacturer’s instructions.

### Flow cytometry

Single-cell suspensions were stained with antibodies in FACS buffer for 40 minutes at 4 °C. Biotinylated antibodies were followed by incubation with fluorochrome-conjugated streptavidin. Dead cells were excluded using the Zombie NIR™ Fixable Viability Kit (BioLegend). In some experiments, cells were fixed in 4% paraformaldehyde (ThermoFisher) for 15 minutes at room temperature before acquisition.

Samples were acquired on an LSR Fortessa (BD Biosciences) using FACS-Diva software and analyzed with FlowJo (v11, BD Biosciences).

The following antibodies were used for flow cytometry:

### Tumor slice supernatant production

67NR tumors were collected in ice-cold Hank’s Balanced Salt Solution (HBSS) (Gibco) and embedded in 4% low-melting point agarose (Invitrogen). The embedded tumors were glued onto the slicing stage of a VT1220S vibratome (Leica Biosystems) and sectioned under the following settings: slice thickness, 250 μm; blade speed,

1.5 mm/s; blade amplitude, 1.25 mm. HBSS served as the cutting medium. Tumor slices were then cultured individually in wells of 24- or 48-well plates using tumor slice culture medium (TSCM) at 37°C with 5% CO₂, as described previously by Nishida-Aoki et al^87^. After 24 hours, supernatants from slices of the same tumor were pooled, centrifuged at maximum speed, filtered three times through 0.22 μm filters (Sigma), and stored at −20°C. Tumor slice supernatants derived from different mice were treated as biological replicates.

### Tumor slice culture medium (TSCM) recipe

| Reagent | Final concentration | Supplier |
| --- | --- | --- |
| William's Medium E |  | Gibco |
| Ascorbic acid | 30μg / ml | Fluorochem |
| Nicotinamide | 12 mM | Sigma |
| HEPES | 10 mM | Gibco |
| Glucose | 5 mg/mL | Gibco |
| Sodium pyruvate | 1x | Gibco |
| GlutaMAX | 1x | Gibco |
| ITS + premix | 1 in 50 | Corning |
| Pen-Strep | 40 IU/mL Pen, 40 µg/mL Strep | Gibco |
| recombinant mouse EGF | 20 ng/mL | Gibco |

### Treatment of tumor slice supernatant

Heat Denaturation - TSS samples were incubated at 100°C for 60 minutes to denature proteins. Fractionation - TSS samples were first centrifuged at 4°C using Amicon Ultra Centrifugal Filter Units (Sigma) with a 3 kDa molecular weight cutoff membrane until 200 μL remained. The >3 kDa fraction was recovered, and blank TSCM was added back to restore the original volume. This process was repeated with 50 kDa cutoff filters (Sigma) to obtain the >50 kDa fraction, followed by volume restoration with blank medium. Lactate Removal - The <3 kDa TSS fraction was incubated for 3 hours at 37°C and 5% CO₂ with varying amounts of lactate oxidase (0, 5, 50, or 500 mU; from *Aerococcus viridans*, Sigma). Post incubation, samples were filtered again through 3 kDa cutoff centrifugal filters (Sigma).

### Conditioned medium production

67NR cells and WT CAFs were cultured in 75 cm² flasks (Greiner) until approximately 60% confluency. The culture medium was then replaced with TSCM. After 24 hours, the conditioned medium was collected, filtered through 0.22 μm filters (Sigma), and stored at −20°C.

### Tumor interstitial fluid

Tumor interstitial fluid was extracted by placing whole tumors in Spin-X filter tubes (Corning) and sequentially centrifuging at 1000 rpm, 2000 rpm, 4000 rpm, and 8000 rpm for 20 minutes at 4 °C. The flow-through was collected, aliquoted, and stored at –20 °C.

### Tumor homogenates

Tumor homogenates were prepared by adding 1 ml RIPA buffer (Sigma) per 100 mg tissue and lysing samples using a Polytron PT 2500 E homogenizer (Kinematica). Lysates were centrifuged at maximum speed for 10 minutes, and the resulting supernatants were collected, aliquoted, and stored at –20 °C.

### Protein quantification by ELISA and Luminex assays

Protein levels were measured using the Quantikine Mouse CCL21/6Ckine ELISA Kit (Bio-techne and the Quantikine Mouse CXCL13/BLC/BCA-1 ELISA Kit (Bio-techne), according to the manufacturer’s instructions. Multiplex quantification of tumor slice supernatants were performed using a custom-designed 20-analyte Luminex assay (Bio-techne), following the manufacturer’s protocol.

### Metabolite quantification by LC-MS

Sample preparation - TSCM, TSS or cell-line conditioned medium was collected and frozen at -80°C. On the day of metabolite extraction, the samples were thawed on ice, vortexed and 140μl of H2O containing 1.5 nmol ^13^C_5_ ^15^N_1_ -valine (Sigma), 150μl methanol (Sigma) and 50μl chloroform (Sigma) was added to 10μl of sample. The samples were spun in a microfuge (4°C, 5 min, 16,000 rpm) and 100μL of the upper aqueous phase containing polar metabolites was extracted for further analysis.

Liquid chromatography - Samples were injected into a Thermo Vanquish UHPLC system (Thermo Fisher Scientific) with a Waters Atlantis Premier BEH z-HILIC (2.1 mm x 100 mm, 1.7 μm particle size) column. A 21 min elution gradient of 5% Solvent A to 95% Solvent B was used, followed by a 2 min wash of 25% of Solvent B and 6 min re-equilibration using the proportion of the initial gradient, where Solvent B was acetonitrile (Optima HPLC grade, Sigma Aldrich) and Solvent A was 12 mM ammonium carbonate in water (Optima HPLC grade, Sigma Aldrich). Other parameters were as follows: flow rate 300 μL/min; column temperature 25°C; injection volume 5 μL; autosampler temperature 4°C.

### Mass spectrometry

Exploris 240 MS was performed with positive/negative polarity switching using an Orbitrap Exploris 240 (Thermo Fisher Scientific) with a HESI II (Heated electrospray ionization) probe. MS parameters were as follows: spray voltage 3.5 kV and 3 kV for positive and negative modes, respectively; probe temperature 320°C; sheath and auxiliary gases were 50 and 10 arbitrary units, respectively; full scan range: 70 to 1000 m/z with settings of normalized AGC target as 100% and resolution as 120,000. Data were recorded using Xcalibur 3.0.63 software (Thermo Fisher Scientific). One-Point mass calibration was performed for both ESI polarities before analysis using Thermo EASY-IC functionality. Parallel reaction monitoring (PRM) acquisition parameters: resolution 60,000; collision energies were set individually in HCD (high-energy collisional dissociation) mode. Data analysis was performed using Compound Discoverer 3.3 software (Thermo Fisher Scientific) employing the embedded metabolomics analysis workflow with mass tolerances set to 3 ppm, peak intensity threshold set at 2e5, and peak rating filter set to 5.

### Lactate measurements

All lactate measurements were conducted following deproteinization using the Deproteinizing Sample Preparation Kit – TCA (Abcam), according to the manufacturer’s instructions. Lactate levels were quantified using the L-Lactate Assay Kit (Colorimetric, Abcam), according to manufacturer’s instructions. For intracellular lactate quantification, 0.5 × 10⁵ in vitro–cultured B cells were washed twice in PBS, homogenized in Lactate Assay Buffer (from the assay kit, Abcam), centrifuged, and the supernatant collected for analysis.

### Lymph node culturing and B cell isolation

Spleens from naïve mice and lymph nodes from naïve or tumor-bearing mice were harvested and mechanically dissociated through a 70 μm filter to create single-cell suspensions. Spleen suspensions underwent red blood cell lysis by incubation in ACK (Gibco) buffer. For transwell migration assays, lymph node cells were cultured for 24 hours in migration medium (RPMI (Gibco) supplemented with 1% BSA (Sigma) and 10 mM HEPES (Gibco)) containing 30 ng/mL BAFF (Bio-techne). B cells were isolated by negative selection using CD43 magnetic beads (Miltenyi) and LD columns following the manufacturer’s protocol.

### Chemotaxis assays

Transwell migration assays were performed using 96-well plates with 5 μm pore inserts (Corning). Fewer than 1 × 10⁵ lymphocytes in 75 μL of migration medium were seeded into the upper chamber. The lower chamber was filled with migration medium, with or without supplementation of 300 ng/mL CCL21 (Peprotech) and/or 300 ng/mL CXCL13 (Peprotech). In some experiments, prior to chemokine addition, the medium was supplemented with sodium lactate (Sigma), lactic acid (Thermo Fisher Scientific), or adjusted to the desired pH using HCl (Sigma). To assess the effect of metabolic inhibitors on B cell migration, glucose-free DMEM (Gibco) (supplemented with 1% BSA (Sigma) and 10 mM HEPES (Gibco)) was used as the base medium. This was further supplemented with either 10 mM glucose (Gibco), 10 mM galactose (Sigma), or 10 mM glucose (Gibco) combined with 1 μM oligomycin (Cell Signaling Technology). For normalization purposes, an equal number of lymphocytes were plated in parallel in a 96-well flat-bottom plate containing the corresponding medium. Cells were allowed to migrate for 3 hours, after which the lower chamber medium (containing migrated cells) and seeding control wells were collected, centrifuged, and stained for flow cytometry to assess cell populations and viability. Samples from both migrated cells and seeding controls were analyzed on an LSR Fortessa (BD Biosciences) using FACS-Diva software (BD Biosciences). Data were processed using FlowJo software (v11, BD Biosciences). The percentage of migrated cells was calculated as: (Number of cells in bottom chamber/Number of cells in seeding control) x 100.

### RNA isolation and RNAseq analysis

A total of 1.5 × 10⁶ magnetically enriched B cells were plated in migration medium and rested for 2 hours. Cells were then treated for 1 hour with either 5 mM sodium lactate (Sigma) or left untreated. Subsequently, 1 μg/mL CXCL13 (Peprotech) and 300 ng/mL CCL21 (Peprotech) were added or omitted, yielding four experimental groups. Three hours after chemokine addition, cells were washed once with ice-cold PBS (Gibco) and lysed in 400 μL Buffer RLT (Qiagen). RNA was extracted using the RNeasy Micro Kit (Qiagen) according to the manufacturer’s instructions. RNA quality and concentration were assessed by spectrophotometric analysis. Library preparation was performed using the RNA Library Prep Kit with polyA enrichment (Watchmaker), according to manufacturer’s instructions. Libraries were sequenced as paired end 100 bp reads, targeting 25 million reads per sample. FASTA files for RNA sequencing were aligned using the GRCm39 reference genome (Ensembl 98). The nf-core/rnaseq pipeline (v3.13.2) was executed, and transcripts were aligned and quantified utilizing the STAR-Salmon method. Downstream RNAseq analysis was conducted in RStudio^88^ (Posit) using the DESeq package. Gene overrepresentation was analyzed via the STRING database (STRING Consortium 2025), and gene set enrichment was performed using the ClusterProfiler and EnrichPlot R packages.

### Glut1 receptor expression

2x10^5^ magnetically enriched B cells were plated in migration medium and rested for 2 hours. Cells were then treated for 30 minutes with either 5 mM sodium lactate (Sigma) or left untreated. Subsequently, 1 μg/mL CXCL13 (Peprotech) and 300 ng/mL CCL21 (Peprotech) were added or omitted, yielding four experimental groups. 30 minutes after chemokine addition, cells were plated and stained with the following antibodies for 30 minutes at 4°C with 1:200 anti-mouse B220 (Clone RA3-6B2; BD Biosciences), 1:200 anti-mouse CD19 (Clone 6D5; BioLegend) and 1:200 anti-mouse Glut1 (Clone EPR3915; Abcam). A Fluorescence Minus One control for the Glut1 antibody was also prepared. Cells were washed once and resuspended in FACS buffer prior to acquisition. Samples were analyzed on an LSR Fortessa (BD Biosciences) using FACS-Diva software (BD Biosciences), and data were processed using FlowJo software (v11, BD Biosciences).

### Actin polymerization assay

Magnetically enriched B cells (2 × 10⁵ per condition) were plated in reduced BSA migration medium (RPMI (Gibco) supplemented with 0.1% BSA (Sigma) and 10 mM HEPES (Gibco)) and rested for 2 hours at 37 °C. Cells were then treated with 5 mM sodium lactate (Sigma) or left untreated for 1 hour. Immediately prior to chemokine stimulation, an aliquot was taken from each well and fixed by adding to three volumes of 4% paraformaldehyde (PFA) (Thermo Fisher Scientific) prepared in cytoskeleton buffer (10 mM MES (Sigma), 138 mM KCl (Sigma), 3 mM MgCl₂ (Sigma), 2 mM EGTA (Sigma) in H₂O). CXCL13 (1 μg/mL) (Peoprotech) and CCL21 (300 ng/mL) (Peprotech) were then added to the remaining cells, and additional aliquots were taken at 10-, 60-, and 180-seconds post-stimulation. Each aliquot was immediately fixed by addition to 4% PFA (Thermo Fisher Scientific) in cytoskeleton buffer as above. Fixed cells were incubated on ice for 20 minutes, centrifuged, and permeabilized for 10 minutes at room temperature in 0.1% Triton X-100 (Sigma) in PBS (Sigma). After centrifugation, cells were blocked for 10 minutes in 2% BSA (Sigma) at 4 °C, centrifuged again, and stained for 20 minutes at room temperature with 1:1000 Alexa 647-conjugated phalloidin (Thermo Fisher Scientific) and 1:200 anti-mouse B220 antibody (clone RA3-6B2; BD Biosciences). Cells were washed twice with PBS (Gibco) containing 0.1% Tween-20 (Sigma) and resuspended in 2% BSA (Sigma) prior to acquisition. Samples were analyzed on an LSR Fortessa (BD Biosciences) using FACS-Diva software (BD Biosciences), and data were processed using FlowJo software (v11, BD Biosciences).

### Mouse and Human histology and immunohistochemistry

Following removal from mice, tumors were fixed in 10% neutral buffered formalin (Scytek Laboratories) overnight at room temperature. In some cases, tumors were bisected to increase surface area. Samples were then transferred to 70% ethanol (Sigma) and processed into formalin-fixed paraffin-embedded (FFPE) blocks using a Tissue-Tek VIP 6 AI processor (Sakura Finetek). Sections (3 μm) were cut and mounted on glass slides (Leica Biosystems).

### 3-plex and 4-plex immunofluorescence

Slides were baked for 1 hour at 60 °C, deparaffinized, and subjected to antigen retrieval by boiling in citrate buffer (pH 6) (Sigma) for 15 minutes. Slides were blocked for 1 hour in blocking buffer (10% normal donkey serum (Sigma), 1% Triton-X (Sigma), 0.5% Fc receptor block (as appropriate, depending on tissue species, BioLegend) in PBS), then incubated for 3 hours in blocking buffer supplemented with 1% anti-mouse Fab fragment (only for mouse tissue, Jackson ImmunoResearch). Overnight incubation with unconjugated primary antibodies was performed at 4 °C. Slides were washed three times in PBS and incubated sequentially with secondary antibodies (2 h, room temperature), washed, and then with conjugated primary antibodies (2 h), followed by a final PBS (Gibco) wash. Slides were mounted with DAPI (1:1000, Thermo Fisher Scientific) in mounting medium (Millipore).

When multiple antibodies raised in the same species were used, staining was performed on the Leica Bond Rx platform (Leica Biosystems). In these cases, 3 μm FFPE sections were baked for 1 hour at 60 °C, blocked with 3% hydrogen peroxide (Millipore) and 0.1% BSA (Sigma), and subjected to antigen retrieval (20 min in Epitope Retrieval Solution 1 or 2, Leica Biosystems) between each staining round. Bond anti-rabbit Polymer (Leica Biosystems) was used as the secondary antibody for rabbit primaries. Slides were counterstained with DAPI (1:2500, Thermo Fisher Scientific, 62248) and mounted with ProLong Gold Antifade reagent (Invitrogen). Whole-slide imaging was performed on an VS120 virtual slide scanner (Olympus) at 20x magnification. For mouse tissue, the following unconjugated primary antibodies and secondary antibodies (including dilutions) were used.

#### Mouse

| Antibody | Clone | Antibody catalogue number | Dilution | Opal | Opal dilution | Secondary antibody | Secondary antibody dilution | Secondary antibody catalogue number |
| --- | --- | --- | --- | --- | --- | --- | --- | --- |
| CD4 | EPR19 514 | ab183685 | 1:750 | Opal 520 | 1:500 | Bond anti-rabbit Polymer | NA | Leica, RE7260-CE |
| CD8 | EPR21 769 | ab217344 | 1:500 | Opal 690 | 1:200 | Bond anti-rabbit Polymer | NA | Leica, RE7260-CE |
| CD19 | EPR23 174-145 | ab245235 | 1:200 | Opal 570 | 1:500 | Bond anti-rabbit Polymer | NA | Leica, RE7260-CE |
| CD19 | EPR23 174-145 | ab245235 | 1:1000 | NA | NA | Alexa Fluor® 647, donkey anti-rabbit | 1:600 | Jackson ImmunoResearch, 711-605-152 |
| CD3 | EPR45 17 | ab134096 | 1:200 | NA | NA | Alexa Fluor® 488, goat anti-rat | 1:600 | Jackson ImmunoResearch, 112-545-003 |

For human tissue, the following unconjugated primary antibodies, secondary antibodies and conjugated primary antibodies were used.

#### Human

| Antibody | Clone | Antibody catalogue number | Dilution | Secondary antibody | Secondary antibody dilution | Secondary antibody catalogue number |
| --- | --- | --- | --- | --- | --- | --- |
| CD20 | EP459Y | Abcam, ab198943, conjugated to Alexa Fluor® 647 | 1:1000 | NA | NA | NA |
| CD21 | EP3093 | Abcam, ab75985 | 1:100 | Alexa Fluor® 488, goat donkey anti-rabbit | 1:600 | Jackson ImmunoResearch, 711-545-152 |

### 6-plex immunofluorescence

3 μm FFPE tumors sections were baked for 1 hour and stained on the Leica Bond Rx platform. Sequential staining was performed as outlined in the Supplementary Tables for the 6-plex panel. Slides were scanned on the PhenoImager HT (formerly Vectra Polaris) in Field mode. Tissue-specific spectral libraries were generated, and spectral unmixing (with autofluorescence removal) was performed using Phenochart™ and InForm® software.

| Position | Antibody | Dilution | Opal | Opal dilution | Antibody catalogue number | Secondary antibody | Antigen retrieval |
| --- | --- | --- | --- | --- | --- | --- | --- |
| 1 | Ly6G | 1:100 | 520 | 1:300 | 87048S | Bond Rabbit Polymer RTU | 20 min Epitope Antigen Retrieval 1 |
| 2 | F4/80 | 1:1000 | 690 | 1:150 | ab300421 | Bond Rabbit Polymer RTU |  |
| 3 | aSMA | 1:250 | 570 | 1:300 | M0851 | Streptavidin-HRP 1:500 cat P0397 Dako |  |
| 4 | CD3 | 1:500 | 480 | 1:100 | ab134096 | Bond Rabbit Polymer RTU |  |
| 5 | CD45 | 1:100 | 620 | 1:200 | 70257S | Bond Rabbit Polymer RTU |  |
| 6 | CD19 | 1:200 | 780 | 1:1000, 1:25 | ab245235 | Bond Rabbit Polymer RTU | 20 min Epitope Antigen Retrieval 2 |

### Immunofluorescence image analysis

Image analysis was performed using QuPath^89^. Cell segmentation was based on DAPI staining, and marker-specific classifiers were developed using channel intensity to identify B cells (CD19 for mouse, CD20 for human), T cells (CD3), CD4⁺ T cells (CD4), and CD8⁺ T cells (CD8). Cell coordinates were exported and analyzed using RStudio (Posit). Digital tissue maps were generated by plotting the X and Y positions of all B cell–classified detections. For clustering analysis, the 50 nearest neighbors within a 50 μm radius of each B cell were analyzed. In mouse samples, tertiary lymphoid structures (TLS) were defined as clusters of more than 50 CD19⁺ cells. In human TNBC tissues, TLS were defined as dense clusters of CD20⁺ and CD3⁺ cells, and mature TLS (mTLS) were defined by the presence of a CD21⁺ follicle.

### Human microarray and RNAseq analyses

Transcriptomic data analyzed in this study were derived from an internal dataset from 124 TNBC patients (Guy’s cohort) and several publicly available datasets (TCGA TNBC^53^, Hammerl TNBC^10^, SCANB TNBC, ISPY-2 breast cancer^55^, merged NSCLC cancer^56^ and Colon cancer^57^). The publicly available datasets were downloaded from the GEO website (National Center for Biotechnology Information). Immune cell deconvolution was performed using the immunedeconv R package, applying the MCP-counter method^41^. B cell score was defined as the MCP Counter B cell score. Survival analyses were conducted in RStudio^88^ using the survival and survminer packages. Cox regression models were used to assess prognostic significance. Gene set enrichment analysis and visualization were carried out using the edgeR, fgsea, ClusterProfiler, and enrichplot packages in R. Aerobic glycolysis cluster assignment was based on hierarchical clustering (Ward’s method, Euclidean distance) of expression data for 12 genes from the Aerobic Glycolysis WikiPathway: *SLC2A1, HK1, GPI, PFKM, ALDOA, TPI1, GAPDH, PGK1, PGAM2, ENO1, PKM2, AND LDHA*. In the Hammerl TNBC cohort, histological B cell scores were extracted from the original publication^10^.

### GeoMx Digital Spatial Profiler Sample Processing

FFPE TNBC tumor tissue sections were processed for spatial transcriptomics using the GeoMx Digital Spatial Profiler (DSP) (nanoString/Bruker Spatial), following the manufacturer’s instructions for FFPE slides. Tissue sections (5 μm) were baked at 60°C for one hour, deparaffinized using xylene, rehydrated, and subjected to antigen retrieval using Antigen Retrieval Solution (Invitrogen). Slides were then incubated with morphology markers (DAPI for nuclei, CD20 (clone EP459Y, ab194970, Abcam) for B cells, and CD3 (clone CD13-12, ab11089, Abcam) for T cells and oligonucleotide-labelled RNA probes targeting the Whole Human Transcriptome. Following scanning, regions of interest (ROIs) were selected and segmented into areas of illumination (AOIs) based on CD20 and CD3 staining patterns: DAPI+CD20+ (B cells), DAPI+CD3+ (T cells), DAPI+CD3+CD20+ (mixed B/T cells), and DAPI+CD3−CD20− (other cells). Photocleavable oligonucleotide probes within each segment were released by sequential UV illumination of the designated AOIs. Library preparation and sequencing were performed according to the manufacturer’s protocols. The GeoMx next-generation sequencing (NGS) pipeline, GeomxTools, was used to convert raw FASTQ files into DCC files containing expression matrices of raw probe counts. These were then converted in R into a GeoMxSet object. AOI quality control was performed by excluding regions that did not meet the following thresholds: total read count greater than twice the number of probes, over 80% trimming and stitching efficiency, more than 75% alignment, less than 50% sequencing saturation, less than 900 non-template control counts, more than 20 nuclei, and a minimum area of 1000 μm². The limit of quantification (LOQ) for each module was calculated using the geometric mean and geometric standard deviation of negative control probes. Specifically, LOQ was defined as the geometric mean multiplied by the geometric standard deviation raised to the power of two. Probes not detected above the LOQ in at least 2.5% of samples were excluded. Gene-level raw counts were calculated as the geometric mean of associated probe counts. Segments were then converted to a SpatialExperiment object using the standR package. Batch correction was performed using the RUV4 method to remove slide-specific effects.

ROIs were manually annotated into five categories based on CD20 and CD3 staining: “Immune Desert”, “Infiltrating”, “Lymphoid Aggregate”, “TLS”, or “Germinal Center”. CXCL13 expression and Aerobic Glycolysis scores were extracted for each AOI. The Aerobic Glycolysis score was defined as the mean expression of 12 genes in the Aerobic Glycolysis pathway (WikiPathways).

### Statistical Analysis

All statistical analyses were performed using GraphPad Prism (V9) and RStudio^88^. A p-value < 0.05 was considered statistically significant.

## Acknowledgements

We thank the members of the Immunity and Cancer laboratory [Francis Crick Institute, FCI, London, UK], the Cancer Bioinformatics [King’s College London (KCL), London, UK], Frederik Igney, Inigo Tirapu, Bino John, Oriana Perez [Boehringer Ingelheim], for critical discussions and comments. We thank Tony Ng [KCL, London, UK] for the 67NR cell line. We thank Marc Hennequart [Université de Namur, Belgium] for the protocol to harvest tumor interstitial fluid. We thank the FCI scientific platforms (Biological Resource Facility, Flow Cytometry, Experimental Histopathology, Light Microscopy, Advanced Sequencing, Metabolomics, Proteomics, Chemical Biology) for expert advice and technical support. Schematics were created with <u>BioRender.com</u>.

## Author contributions

Conceptualization: V.B., A.G., and D.P.C.

Methodology: V.B., E.A., P.D., M.S.H., A.C., J.Q., M.S, and J.M.

Investigation: V.B., E.A., and P.D.

Resources: A.G., F.L., J.R., C.G., L.A., V.P., and D.P.C.

Visualization: V.B.

Funding acquisition: A.G., and D.P.C.

Supervision: A.G. and D.P.C.

Writing-original draft: V.B., A.G., and D.P.C. Writing-review and editing: All authors.

---

**Supp figure 1.**
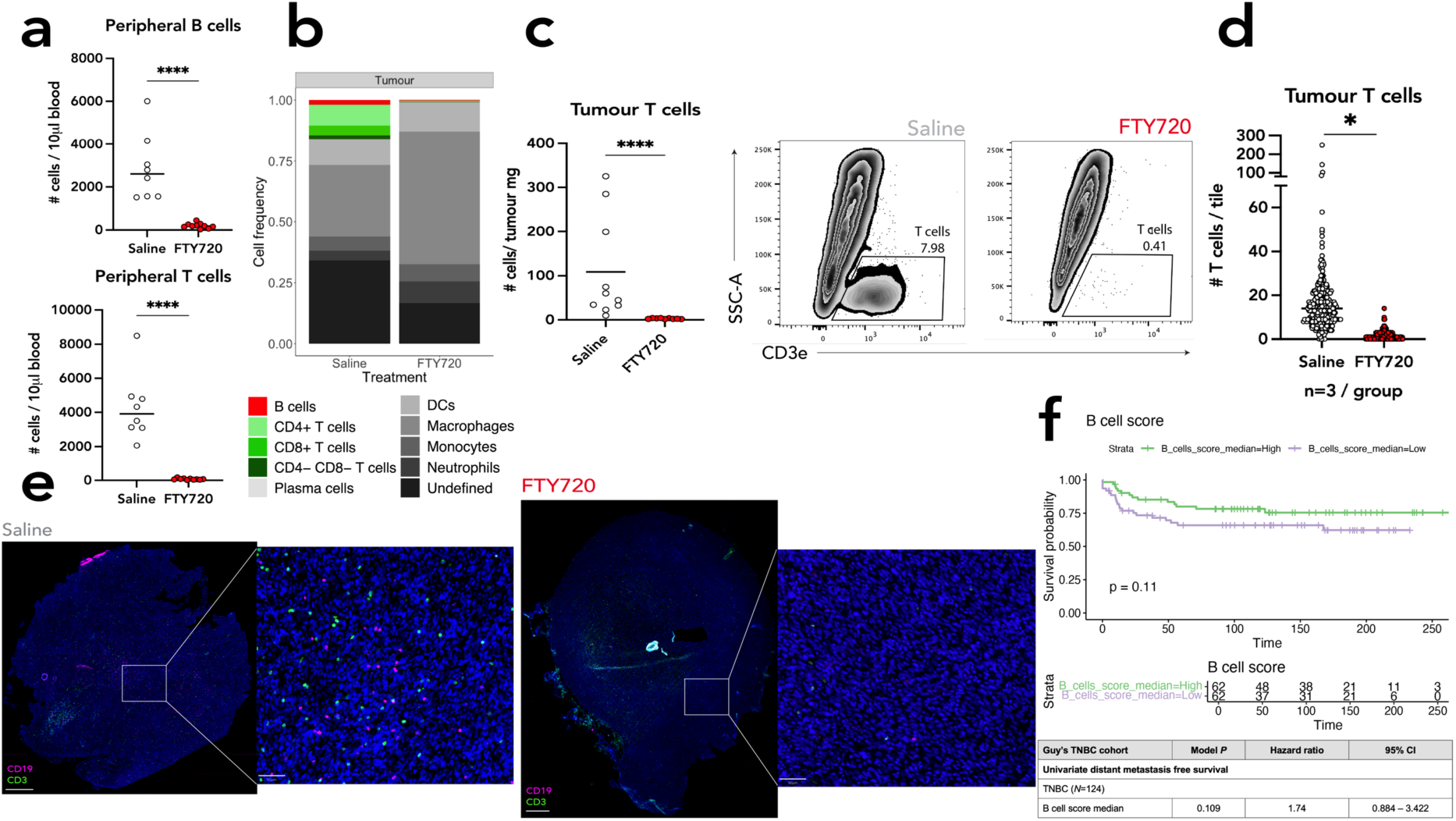
Characterization of B cell abundance and immune modulation in murine breast tumors models and clinical correlates. **(a)** *Top*: Quantification of circulating B cells in 67NR tumor-bearing mice treated with saline or FTY720 (cells per 10 *μ* L of blood, day 7). *Bottom*: Quantification of circulating T cells in the same mice (day 7). **(b)** Immune composition of the 67NR tumor microenvironment in saline- or FTY720-treated mice at day 14, with B cells in red and T cells in shades of green. **(c)** *Left*: Quantification of tumor-infiltrating T cells (cells per mg tumor) in saline- or FTY720- treated mice (day 14). *Right*: Representative gating strategy for tumor-infiltrating T cells (gated on CD45⁺). **(d)** Quantification of tumor-infiltrating T cells by immunofluorescence (cells per tile, day 14). **(e)** Immunofluorescence image and inset of 67NR saline- or FTY720-treated tumors stained for B cells (CD19, magenta) and T cells (CD3, green). Scale bars: 400 μm, 50 μm. **(f)** *Top*: Kaplan–Meier plot showing distant metastasis occurrence in patients stratified by high vs. low B cell score. *Bottom*: Univariate Cox regression model comparing B cell score groups. In panels (c), (d) and (f), each dot represents an individual mouse [(c) *n* = 4, (d) Saline *n* = 8, FTY720 *n* = 10, (f) Saline *n* = 10, FTY720 *n* = 10]. In panel (g), each dot represents an individual tile and the tiles from individual mice have been combined [(g) Saline *n*=3, FTY720 *n*=3]. Bars represent the mean. Data in (c) are from one independent experiment. Data in (d) and (g) are from the same two independent experiments. \*\**P* ≤ 0.01; \*\*\*\**P* ≤ 0.0001 [paired Friedman test with Dunn’s multiple comparison correction in (c); Mann-Whitney U test in (d), (f); nested Student’s *t* test in (g)].

**Supp figure 2.**
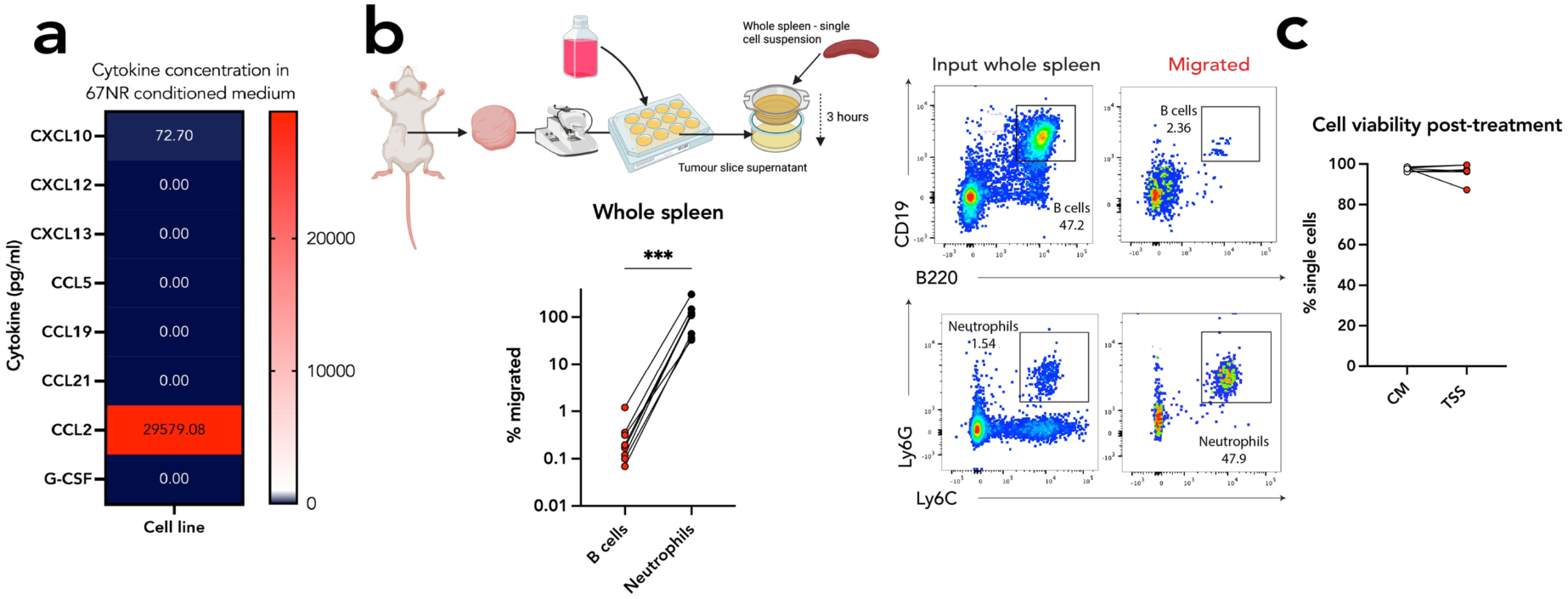
Tumor secreted factors disrupt lymphocyte chemotaxis by inhibiting chemokine responsiveness. **(a)** Heatmap showing concentrations (pg/mL) of selected cytokines and chemokines in 67NR- conditioned medium. **(b)** *Top*: Schematic of transwell chemotaxis assay testing migration of whole spleen immune populations toward blank tumor slice supernatant (TSS). *Bottom*: Paired quantification of spleen cell migration toward blank TSS. Right: Representative gating strategy for input and migrated spleen B cells and neutrophils. **(c)** Viability of lymphocytes (% of single cells) after 3-hour incubation in control medium (CM) or TSS. Each symbol represents an individual mouse [(b) n = 8; (c) n = 5]. Bars represent the mean. Data in (b) are from 2 independent experiments. Data in (c) are from one independent experiment. \*\*\**P* ≤ 0.001. [Paired Wilcoxon test in (b); paired Wilcoxon test in (c)].

**Supp figure 3.**
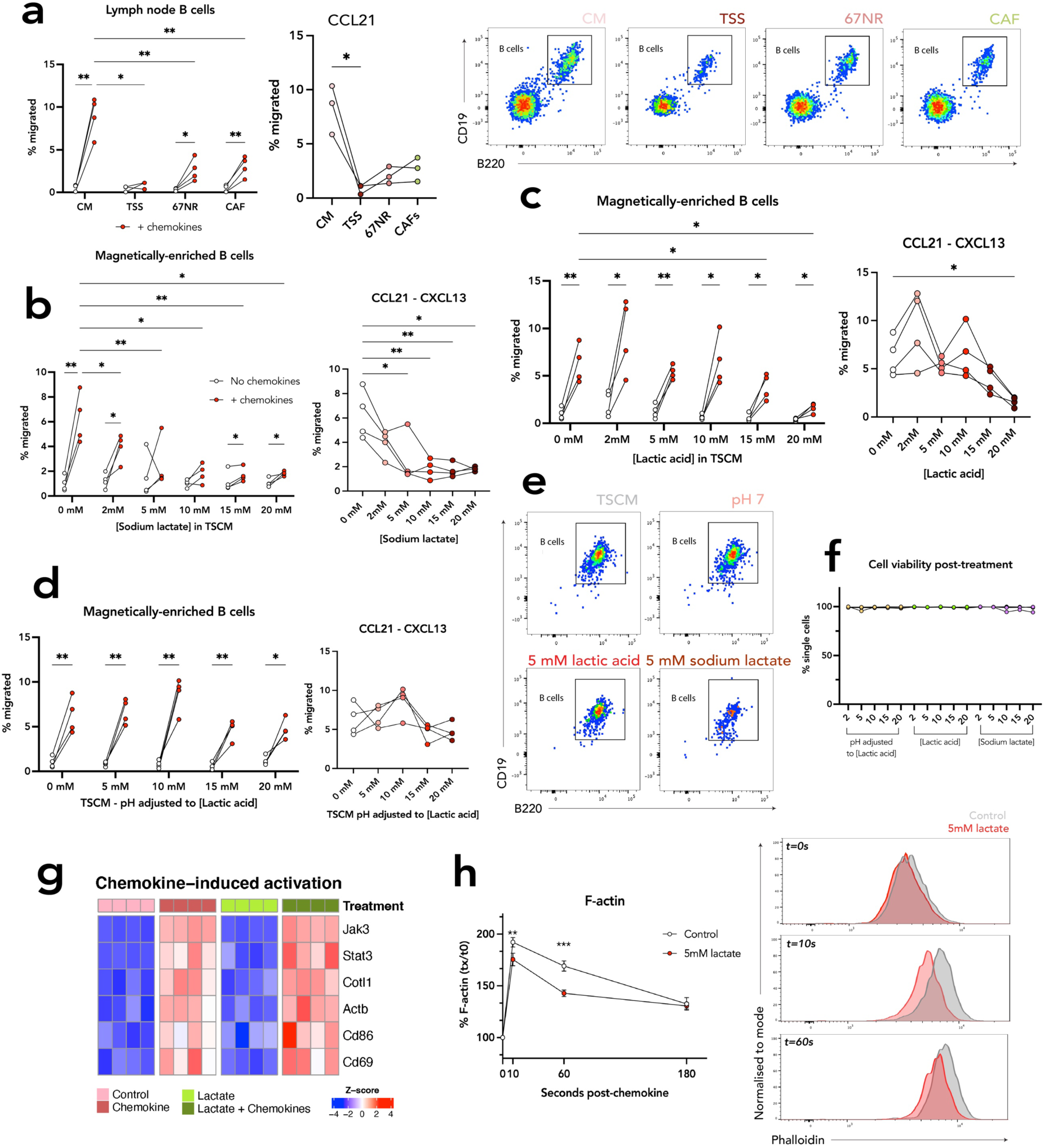
Lactate and tumor-derived factors inhibit chemokine-mediated B cell migration via metabolic and cytoskeletal reprogramming. **(a)** *Left:* Quantification of B cell migration towards blank CM, TSS, 67NR- or CAF-conditioned medium, with or without chemokine supplementation. *Middle:* Paired migration quantification of purified B cells towards CCL21-supplemented CM, TSS, 67NR, or CAF-conditioned medium. *Right:* Representative gating strategy for migrated B cells. **(b)** *Left:* B cell migration in CM with increasing concentrations of sodium lactate, with or without chemokine supplementation. *Right:* Paired migration quantification of purified B cells towards chemokine-supplemented CM with increasing sodium lactate concentrations. **(c)** *Left:* B cell migration in CM with increasing concentrations of lactic acid, with or without chemokine supplementation. *Right:* Paired migration quantification of purified B cells towards chemokine-supplemented CM with increasing lactic acid concentrations. **(d)** *Left:* B cell migration in CM pH-matched with HCl to mimic increasing concentrations of lactic acid, with or without chemokine supplementation. *Right:* Paired migration quantification of purified B cells towards chemokine-supplemented, pH-adjusted CM. **(e)** Representative gating strategy for migrated B cells towards chemokine-supplemented blank CM, 5 mM sodium lactate, 5 mM lactic acid, or pH 7 CM. **(f)** *Top:* B cell viability after 3-hour incubation in CM containing increasing concentrations of sodium lactate, lactic acid, or decreasing pH. **(g)** Scaled heatmap showing expression of chemokine-induced activation genes across B cell treatment conditions. **(h)** *Left:* Percentage increase in F-actin polymerization following chemokine stimulation in B cells pretreated or not with 5 mM lactate. *Right:* Phalloidin fluorescence intensity at defined time points post-chemokine stimulation in B cells with or without lactate pretreatment. Except when otherwise stated, each symbol represents an individual mouse [(a) n = 4; (b) n =4; (c) n = 4; (d) n = 4; (f) n =4]. In (h), the dot represents the mean, and the bar represents the SEM [(i) n = 4]. Data in (a) are from one independent experiment. Data in (b), (c), (d) and (f) are from the same independent experiment. Data in (h) is representative of two experiments. \**P* ≤ 0.05; \*\**P* ≤ 0.01; \*\*\**P* ≤ 0.001. [Paired mixed effects analysis in left (a), paired Friedman test with Dunn’s multiple comparison correction in middle (e); paired two-way ANOVA in left (b), paired Friedman test in right (e); paired two-way ANOVA in left (c), paired Friedman test in right (c); paired two-way ANOVA in left (d), paired Friedman test in right (d); paired Friedman test with Dunn’s multiple comparison correction in (f); paired two-way ANOVA in (h)].

**Supp figure 4.**
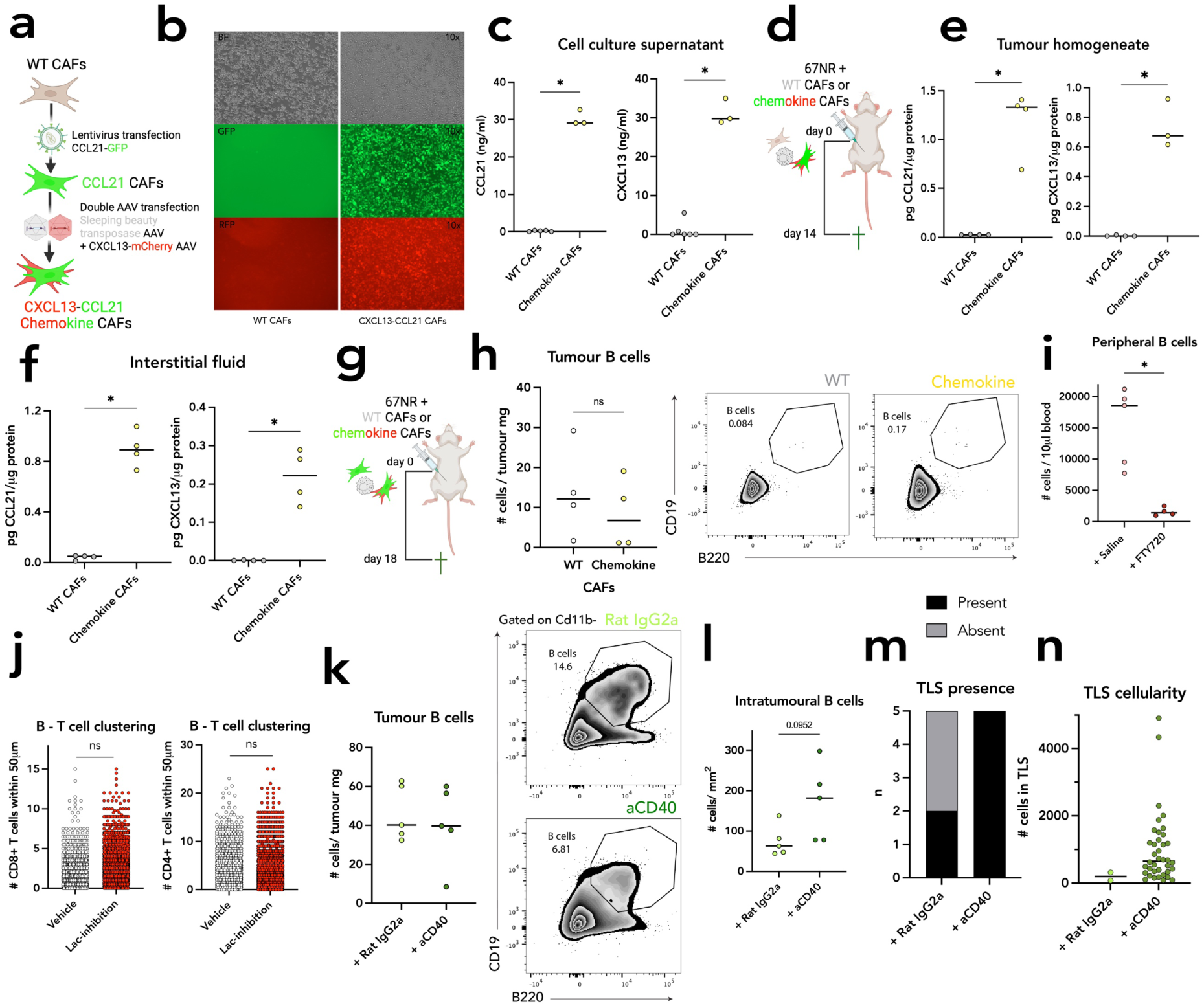
Restriction of lactate production enhances chemokine-driven lymphocyte recruitment and TLS formation. **(a)** Schematic of chemokine CAF generation protocol. **(b)** Live-cell imaging of WT and chemokine CAFs showing GFP and mCherry fluorescence. **(c)** Chemokine quantification by ELISA in supernatants from WT and chemokine CAFs (Left: CCL21; Right: CXCL13). **(d)** Schematic of in vivo co-injection of 67NR with WT or chemokine CAFs for tumors harvest. **(e)** Chemokine quantification by ELISA in tumors homogenates from 67NR+WT CAF or 67NR+chemokine CAF tumors (Left: CCL21; Right: CXCL13). **(f)** Chemokine quantification by ELISA in tumor interstitial fluid from 67NR+WT CAF or 67NR+chemokine CAF tumors (Left: CCL21; Right: CXCL13). **(g)** Schematic of tumor harvest at day 18 post co-injection of 67NR and CAFs. **(h)** *Left:* Quantification of tumor-infiltrating B cells by flow cytometry (cells per mg of tumor) on day 14. *Right:* Representative gating of CD45⁺ tumor-infiltrating B cells (day 18). **(i)** Quantification of circulating B cells in mice treated with lactate inhibition and either saline or FTY720 (cells per 10 µL blood, day 11). **(j)** Quantification of B-T cell clustering by immunofluorescence in 67NR+chemokine CAF tumors ± lactate inhibition (Left: B–CD4⁺; Right: B–CD8⁺; shown as number of cells within 50 µm per B cell). **(k)** *Left:* Flow cytometric quantification of tumor-infiltrating B cells in mice treated with lactate inhibition and either isotype control or aCD40 (cells per mg, day 14). *Right:* Gating of B cells on Cd11b^-^ cells. **(l)** B cell density (cells per mm²) in tumor sections by immunofluorescence under conditions in (q). **(m)** Contingency plot showing TLS presence or absence under isotype or aCD40 treatment. **(n)** Quantification of TLS cellularity (per TLS) in mice treated with lactate inhibition and either isotype or aCD40. Except when otherwise stated, each symbol represents an individual mouse [(e) and (f) 67NR+WT n = 4, 67NR+chemokine CAF n = 4; (h) 67NR+WT n = 4, 67NR+chemokine CAF n = 4; (i) +Saline n = 5, +FTY720 n = 4; (k) +Rat IgG2a n = 5; +aCD40 n = 5; (l) +Rat IgG2a n = 5; +aCD40 n = 5]. In (c), each dot represents a technical replicate [(c) WT n = 6, Chemokine n = 3]. In (j), each dot represents an individual B cell and the B cells from individual mice have been combined [(j) Vehicle n = 6, Lac-inhibition n = 6]. In (n), each dot represents an individual TLS and TLS from individual mice have been combined [(n) +Rat IgG2a n=2, +aCD40 n=5]. Bars represent the mean. Data in (c), (h) and (i) are from one independent experiment. Data in (e) and (f) are from the same independent experiment. Data in (j) are from two independent experiments. Data in (k), (l) and (n) are from the same independent experiment. \**P* ≤ 0.05; \*\**P* ≤ 0.01; \*\*\**P* ≤ 0.001. [Mann-Whitney U test in (c), (e), (f), (h), (i), (k) and (l); nested Student’s t test in (j) and (n)].

**Supp figure 5.**
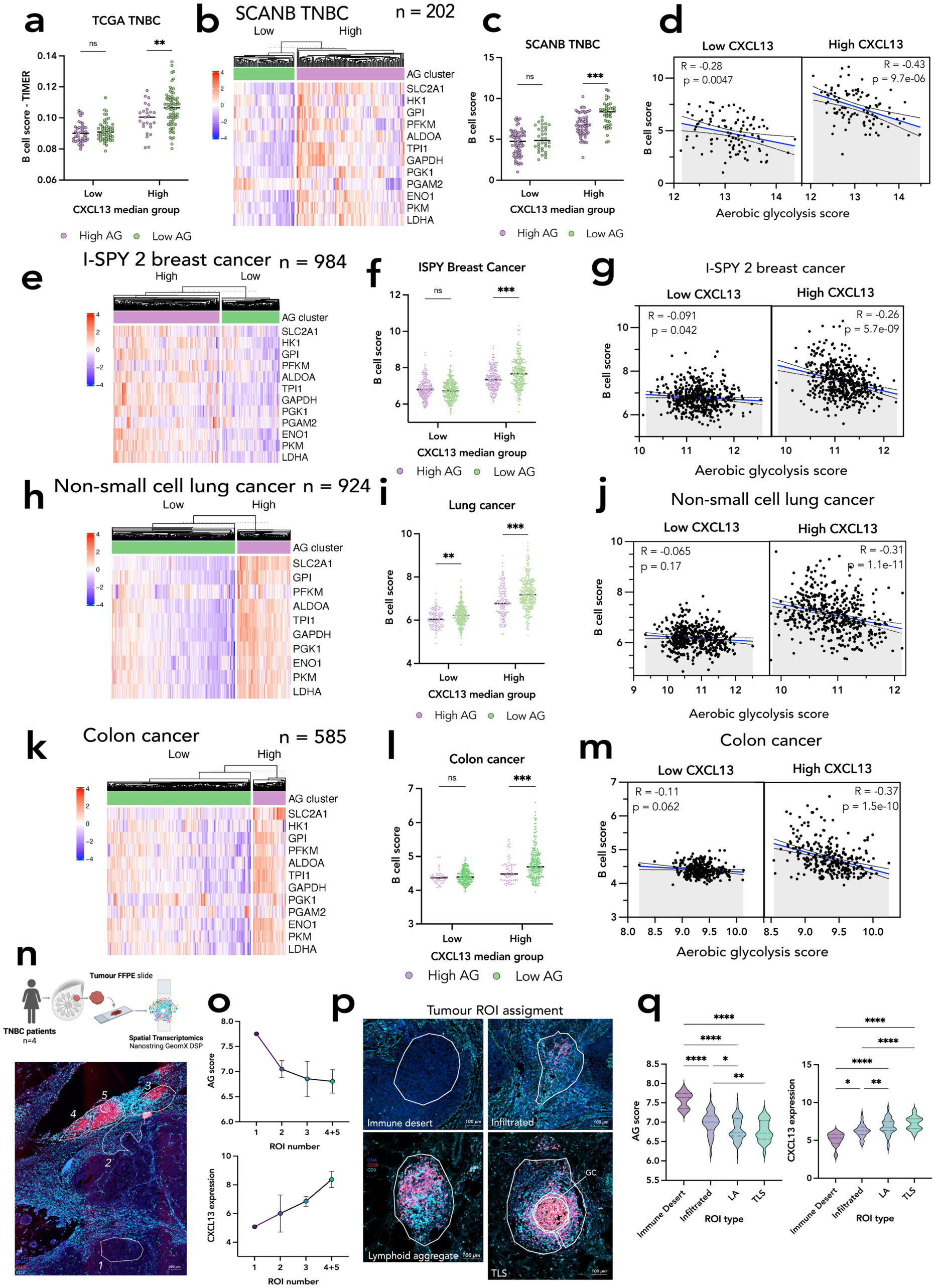
Aerobic glycolysis is associated with reduced B cell infiltration in CXCL13-high tumors across multiple cancer types and spatial contexts. **(a)** B cell scores derived from TIMER immune deconvolution across four Aerobic Glycolysis (AG) and CXCL13 expression groups in the TCGA TNBC cohort. **(b)** Heatmap showing scaled expression of 12 AG-related genes in SCANB TNBC patients (*n* = 202) grouped by hierarchical clustering. **(c)** MCP B cell scores across AG/CXCL13 groups in the SCANB TNBC cohort. **(d)** Correlation between AG score and MCP B cell score in SCANB TNBC patients, stratified by CXCL13 expression (high vs. low, median split). **(e)** Heatmap of scaled AG gene expression in ISPY2 TNBC patients (*n* = 984), grouped by hierarchical clustering. **(f)** MCP B cell scores across AG/CXCL13 groups in the ISPY2 TNBC cohort. **(g)** Correlation between AG score and MCP B cell score in ISPY2 TNBC patients, split by CXCL13 median expression. **(h)** Heatmap of scaled AG gene expression in NSCLC patients (*n* = 984), split by hierarchical clustering. **(i)** MCP B cell scores across AG/CXCL13 groups in the NSCLC cohort. **(j)** Correlation between AG score and MCP B cell score in NSCLC patients stratified by CXCL13 expression. **(k)** Heatmap of scaled AG gene expression in colon cancer patients (*n* = 585), grouped by hierarchical clustering. **(l)** MCP B cell scores across AG/CXCL13 groups in the colon cancer cohort. **(m)** Correlation between AG score and MCP B cell score in colon cancer patients stratified by CXCL13 expression. **(n)** *Top:* Schematic of spatial transcriptomics using the GeoMx DSP platform on four TNBC patient tumors from the Tianjin cohort. *Bottom:* Immunofluorescence image showing regions of interest (ROIs) selected for transcriptomic profiling, with staining for B cells (CD20, red) and T cells (CD3, cyan). **(o)** Aerobic Glycolysis scores (*top*) and CXCL13 expression (*bottom*) in merged areas of illumination (AOIs), ordered by increasing B cell infiltration from the ROIs shown in (s). **(p)** Representative GeoMx DSP immunofluorescence images for each ROI type, stained for B cells (CD20, red) and T cells (CD3, cyan). **(q)** Aerobic Glycolysis scores (*left*) and CXCL13 expression (*right*) across the different ROI categories defined in the spatial transcriptomics dataset. Except when otherwise stated, each symbol represents an individual patient [(a) TCGA TNBC: Low CXCL13 High AG n = 45, Low CXCL13 Low AG n = 51, High CXCL13 High AG n = 25, High CXCL13 Low AG n = 70; (c) SCANB TNBC: Low CXCL13 High AG n = 67, Low CXCL13 Low AG n = 34, High CXCL13 High AG n = 62, High CXCL13 Low AG n = 39; (d) SCANB TNBC: Low CXCL13 n = 101, High CXCL13 n = 101; (f) ISPY2 TNBC: Low CXCL13 High AG n = 220, Low CXCL13 Low AG n = 285, High CXCL13 High AG n = 206, High CXCL13 Low AG n = 271; (g) ISPY2 TNBC: Low CXCL13 n = 505, High CXCL13 n = 477; (i) NSCLC: Low CXCL13 High AG n = 128, Low CXCL13 Low AG n = 334, High CXCL13 High AG n = 151, High CXCL13 Low AG n = 311; (j) NSCLC: Low CXCL13 n = 462, High CXCL13 n = 462; (l) Colon cancer: Low CXCL13 High AG n = 46, Low CXCL13 Low AG n = 246, High CXCL13 High AG n = 62, High CXCL13 Low AG n = 229; (m) Colon cancer: Low CXCL13 n = 292, High CXCL13 n = 291]. Bars represent the mean. In (o), each symbol represents the mean of the AOIs for that ROI, bars are the SEM. In (q), the violin plots represent data from individual AOIs within ROIs assigned each category from n = 4 TNBC patient tumors. Data in (o) and (q) are from the same independent experiment. *P ≤ 0.05; **P ≤ 0.01; ***P ≤ 0.001; **** P ≤ 0.0001. [Two-way ANOVA in (a), (c), (f), (i) and (l), only results within CXCL13 groups are shown; Pearson’s correlation test in (d), (g), (j) and (m); Kruskal-Wallis tests in (q)].

